# Multiomics Characterization Identifies S1P-secreting USS^High^ Skeletal Muscle Stem Cells as Essential Drivers of Niche Remodeling and Muscle Regeneration

**DOI:** 10.64898/2026.09.18.752516

**Authors:** Xingyuan Liu, Yingkun Zhang, Yang Li, Yi Jin, Yulong Qiao, Chuhan Li, Siyuan Wang, Sin Man Lam, Guanghou Shui, Catherine C.L. Wong, Hao Sun, Shihuan Kuang, Huating Wang

## Abstract

Acutely induced cellular senescence is increasingly recognized as vital for tissue repair but the precise roles of senescent cells in skeletal muscle regeneration remain paradoxical. Here, using single-cell RNA sequencing and a Unified Senescence Score (USS) methodology, we identify a distinct USS^High^ (USS^Hi^) muscle stem cell (MuSC) subpopulation emerging in early phase of acute injury induced muscle regeneration. Isolation via the surface marker MHCII enables characterization of these cells as exhibiting canonical senescent-like features. Further characterization through integrated transcriptomics, proteomics, and secretomics reveals their highly pro-inflammatory secretory capacity. Moreover, lipidomic profiling uncovers that the USS^Hi^ MuSCs undergo profound sphingolipid metabolic remodeling to generate upregulated bioactive lipid Sphingosine-1-phosphate (S1P). *In vivo* specific ablation of this subpopulation using a *Pax7-CreERT; Ccl2-LSL-DTA* genetic strategy significantly impaired muscle regeneration. Furthermore, spatial transcriptomics and cell-cell communication analyses revealed that USS^Hi^ MuSCs act as critical niche modulators, directing crosstalk with multiple cell types. Specifically, S1P is essential for mediating direct USS^Hi^ interaction with endothelial cells and USS^Lo^ cells to promote endothelial angiogenesis and MuSC expansion in the regenerating muscle. Crucially, exogenous S1P rescues the regenerative deficits caused by USS^Hi^ MuSC ablation. Altogether our findings define a senescent-like MuSC population emerging in acute injury induced muscle regeneration and provide definitive evidence to demonstrate the positive role of these cells in enhancing muscle repair via their robust secretory phenotypes and inter-cellular communicative capacity.

## Introduction

Skeletal muscle accounts for 30–40% of human body mass and is indispensable for both systemic metabolic homeostasis and locomotion^1^. This highly plastic tissue exhibits a robust capacity for regeneration following acute injury, a process fundamentally driven by the activation and proliferation of muscle stem cells (MuSCs). Under homeostatic conditions, MuSCs are maintained in a quiescent state beneath the basal lamina, uniquely labeled by the expression of paired box 7 (PAX7)^2^. Upon extrinsic stimuli, such as injury, these stem cells exit quiescence, enter the cell cycle, and transition into proliferating myoblasts driven by key transcription factor MYOD induction; the myoblasts ultimately differentiate into myocytes with upregulated Myogenin expression and fuse to repair damaged myofibers^3,4^. This regenerative cascade is tightly coordinated by a dynamic niche microenvironment comprising immune cells, fibro-adipogenic progenitors (FAPs), and endothelial cells (ECs) etc., which collectively orchestrate tissue repair through complex cellular crosstalk^5^. For instance, immediately following injury, the rapid infiltration of neutrophils clears necrotic debris and recruits pro-inflammatory macrophages, which secrete factors to stimulate MuSCs activation and proliferation. A subsequent pivotal shift to anti-inflammatory macrophages resolves inflammation and drives the differentiation and fusion of myoblasts^6^. Parallel to this immune dynamic, angiogenesis distinctly precedes myogenesis to restore the essential vascular niche. ECs actively regulate this process by providing metabolic substrates and secreting angiocrine factors that govern MuSC activation and proliferation^7^.

Recently, cellular senescence has emerged as a critical, albeit highly complex, stress-induced regulatory program influencing this regenerative niche. Senescence is defined as a state of stable, virtually irreversible cell-cycle arrest, typically orchestrated by the activation of the p53/p21 and p16^INK4a^/pRB tumor suppressor pathways^8^. Beyond growth arrest, senescent cells undergo marked phenotypic alterations, most notably the acquisition of a senescence-associated secretory phenotype (SASP). The SASPs involve the robust secretion of pro-inflammatory cytokines, chemokines, growth factors, and extracellular matrix-modifying enzymes^9,10^. The chronic accumulation of senescent cells is widely recognized as a primary driver of tissue aging and age-related degenerative pathologies^11^. Our recent study mapped the first cellular senescence atlas in aging human muscle and demonstrated senescent MuSCs contribute to the deteriorating aging niche via SASP production^12^. Conversely, transient senescence arising in developmental or regenerating tissues is thought to exert a beneficial, programmed mechanism during embryonic development and specific tissue repair processes^13^. For example, the transient induction of senescence within ECs and fibroblasts expedites cutaneous wound healing by secreting SASP factor PDGF-AA to promote angiogenesis and myofibroblast differentiation^14^.

In the context of skeletal muscle regeneration, however, the precise functional role of cellular senescence remains controversial, yielding contradictory findings across recent literature. A subset of pioneering studies supports a transient, pro-regenerative role for senescence^15–17^. Evidence indicates that during acute injury in young mice, a transient population of senescent cells (particularly FAPs and macrophages) emerges locally within the tissue. Through the timely secretion of specific SASP factors, these cells are thought to enhance cellular plasticity, facilitate the phagocytic clearance of necrotic debris, and promote myoblast differentiation. Global selective clearance of the senescent cells via senolytic drugs such as ABT-263 or Dasatinib and Quercetin (D+Q) markedly impairs muscle regeneration ^18,19^. Unexpectedly, a recent study suggests that senescence exerts profoundly detrimental effects on muscle repair^20^. It is shown that senescent cells, including myeloid cells, FAPs and MuSCs contribute to the establishment of an aged-like, pro-inflammatory, and pro-fibrotic niche that blunts MuSC proliferation and impedes optimal tissue repair. In both aged and young mouse models, D+Q treatment or genetic models have been shown to reduce fibrosis, lower inflammation, and enhance regenerative outcomes, arguing that senescence inherently restricts muscle recovery. We believe a few factors drive these conflicting findings. First, emerging studies including ours highlight the vast heterogeneity of senescent cells in terms of their cell type specific markers and the highly variable SASP signatures^21,22^. Reliance on p16^INK4a^ expression alone is thus insufficient as a universal marker for the identification and ablation of senescent cells. The previous studies predominantly relying on global senescent cells clearance strategies like senolytic drugs or *p16-3MR* mice may therefore conflate the distinct roles of each type of senescent cells^18,20,23^. Additionally, the *p16-3MR* ablation model has been shown to exhibit inefficient clearance of senescent cells, confounding data interpretation^24^. We reason that to definitively resolve the functional impact of cellular senescence in muscle repair, it is critical to define, isolate and characterize specific types of senescent cells individually and a genetic approach adequately removing the senescent cells is a prerequisite for functional elucidation.

Here we focus our study on MuSCs aiming to define and characterize the senescent phenotype of MuSCs ultimately to clarify their exact functions and mechanism in acute injury induced muscle regeneration. Utilizing a computational scoring strategy, Unified Senescence Score (USS), alongside a targeted genetic ablation model, we identify a special USS^High^ (USS^Hi^) USS^Hi^ MuSC subpopulation with senescence features and demonstrate that these cells are indispensable orchestrators of skeletal muscle repair rather than retarders. High-resolution spatial transcriptomics and scRNA-seq based cell-cell communication analysis reveal that these cells function as vital paracrine signaling hubs in the muscle injury microenvironment. Multi-omics characterization further uncovers that USS^Hi^ MuSCs undergo profound sphingolipid metabolic remodeling to secrete the bioactive lipid Sphingosine-1-phosphate (S1P), which mediates the crosstalk with ECs and USS^Low^ (USS^Lo^) MuSCs to drive localized angiogenesis and MuSC expansion (Fig. 1A). Together, these findings demonstrate the essential beneficial role of USS^Hi^ MuSCs in the skeletal muscle regeneration process and highlight the regenerative importance of lipid-mediated signaling.

**Figure 1.**
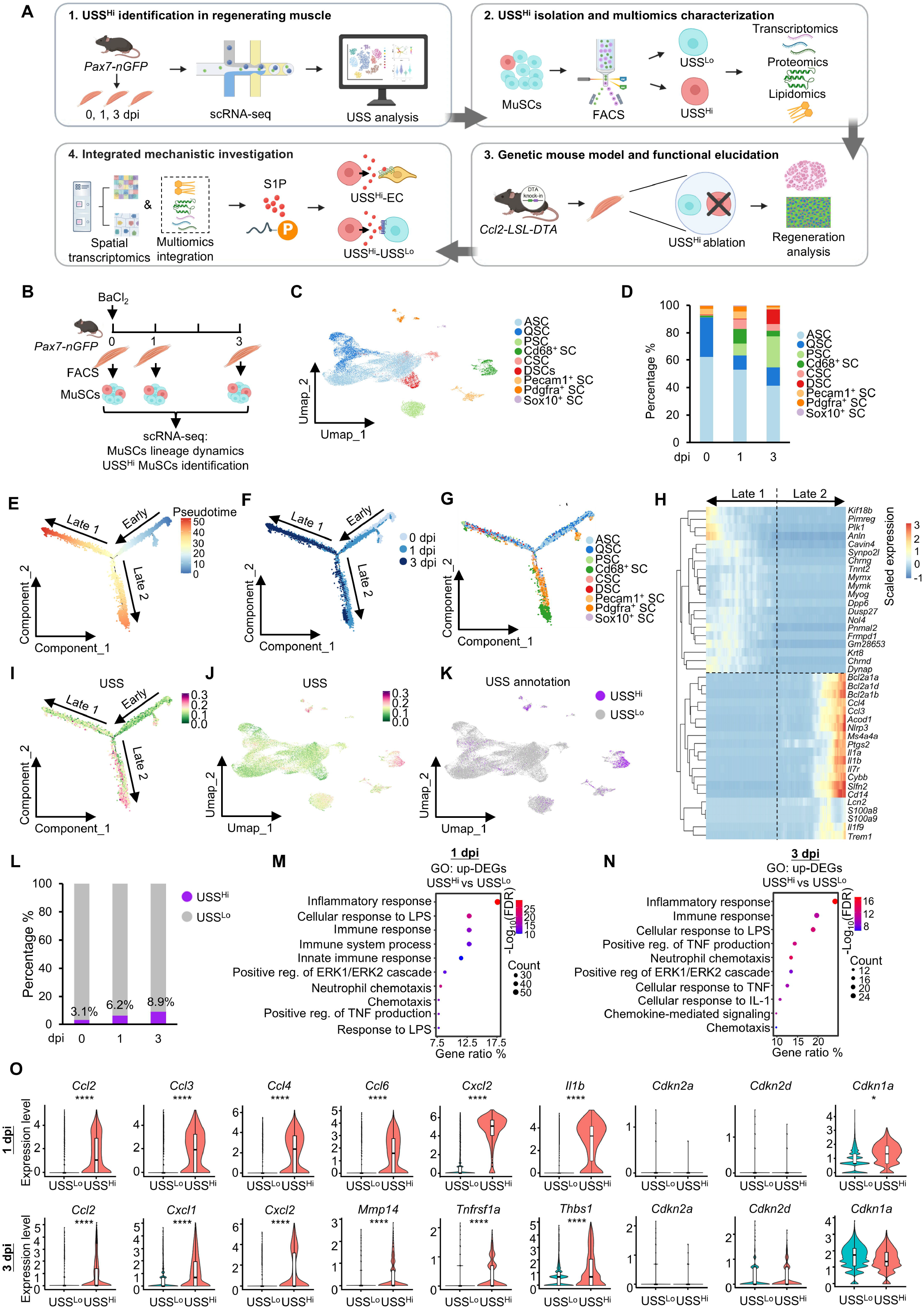
Single-cell mapping delineates MuSC activation dynamics and identifies USS^Hi^ subpopulation in regenerating skeletal muscle. **(A)** Schematic overview of the study workflow. **(B)** Schematic illustration of scRNA-seq mapping of MuSC activation dynamics in acute injury induced muscle regeneration: acute injury was induced by BaCl_2_ injection into the TA of 3-month-old *Pax7-nGFP* mice, and MuSCs were isolated by FACS at 0, 1 and 3 dpi of the regeneration course for scRNA-seq analysis. **(C)** UMAP projection of all the above captured MuSCs colored by the nine annotated subclusters. **(D)** Stacked bar chart showing the relative proportion of each of the above identified cell clusters at 0, 1, and 3 dpi. **(E-F)** Pseudotime trajectory analysis revealing a bifurcated progression from the Early state toward the Late 1 and Late 2 fate branches, colored by pseudotime (E) and by time point (F). **(G)** Dynamics of the above MuSC subpopulation proportions along the pseudotime of each fate branch. **(H)** Hierarchical clustering heatmap of the top 20 DEGs distinguishing the Late 1 and 2 branches. **(I-J)** USS projected onto the pseudotime trajectory (I) and the UMAP space (J). **(K)** UMAP projection showing the segregation of USS^Hi^ and USS^Lo^ MuSCs. **(L)** Quantification of the USS^Hi^ MuSC fraction across the regeneration timeline at 0, 1 and 3 dpi. **(M-N)** GO enrichment analysis of upregulated DEGs in USS^Hi^ vs. USS^Lo^ MuSCs at 1 dpi (M) and 3 dpi (N). **(O)** Violin plots showing expression of inflammatory SASPs and commonly used senescence markers *Cdkn2a* (p16^INK4a^), *Cdkn2d* (p19) and *Cdkn1a* (p21) in USS^Hi^ vs. USS^Lo^ MuSCs at 1 and 3 dpi. Unpaired two-sided Student’s t-test was used to calculate the statistical significance: \**p* <0.05, \*\**p* <0.01, \*\*\**p* <0.001, \*\*\*\**p*<0.0001, *ns* = no significance.

## Results

### Single-cell mapping delineates MuSC activation dynamics and identifies USS^Hi^ subpopulation in regenerating skeletal muscle

To elucidate the dynamic senescence landscape of MuSCs during acute injury induced muscle regeneration, we generated high-resolution scRNA-seq datasets from freshly isolated MuSCs across distinct regenerative stages (Fig. 1A-B) instead of using the published whole muscle scRNA-seq datasets^23^ that may not capture the entire population of MuSCs. Acute injury was induced in the tibialis anterior (TA) muscles of 3-month-old *Pax7-nGFP* mice via a standard intramuscular injection of BaCl_2_ ^25^ (Fig. 1B). Shortly following the injury, the necrosis of damaged myofibers triggered an inflammatory response characterized by the infiltration of neutrophils and activated macrophages, which established an inflammatory microenvironment. Quiescent MuSCs were subsequently activated within 1 day post injury (dpi), undergoing extensive proliferation into myoblasts until around 3 dpi, which then initiate differentiation to repair damaged myofibers in later stage of the regeneration (5 dpi). To capture the lineage progression dynamics, MuSCs were isolated by fluorescence-activated cell sorting (FACS) from the injured muscle at 0, 1, and 3 dpi and subjected to scRNA-seq (Fig. 1B and Suppl. Dataset 1). Following quality control and filtering, a total of 18,083, 15,587 and 14,920 qualified cells were captured at each time point. Uniform Manifold Approximation and Projection (UMAP) dimensionality reduction revealed a total of 9 unique subclusters (Fig. 1C), which were annotated according to their canonical gene expression profiles (Suppl. Fig 1A). Specifically, quiescent MuSCs (QSC) expressed high level of *Pax7*, while activated MuSCs (ASC) co-expressed *Pax7* and *Myod1*. Proliferating MuSCs (PSC) or myoblasts were marked by *Pax7* and *Myod1* along with the cell proliferation markers *Mki67* and *Top2a*. Interestingly, we found four subpopulations expressing MuSC marker (*Pax7*, *Myod1*) simultaneously with high levels of *Cd68*, *Pdgfra, Pecam1* or *Sox10* respectively. Consistent with previous investigation^26^, a subpopulation representing the intermediate, proliferating phase bridging stem cell activation and terminal differentiation, termed committed MuSCs (CSC) was also identified and characterized by the co-expression of the differentiation marker *Myog* and proliferation markers. Differentiated MuSCs (DSC) robustly expressed *Myog* and mature muscle marker *Myh3.* Assessment of the cellular composition across the regenerating timeline highlighted the sequential state transition of MuSCs (Fig. 1D): the 0 dpi pool was dominated by QSC and ASC, with ASCs likely reflecting a basal activation triggered by the isolation procedure^27,28^. Following injury (1 dpi), the percentages of QSC and ASC dropped rapidly as cells mobilized into the cell cycle, leading to an influx of PSC and a transitional CSC. By 3 dpi, the regenerative program progressed further: PSC continued to rise, whereas CSC transitioned into a growing population of DSC. These temporal dynamics perfectly mirror the classical cascade of MuSC activation, proliferation, and terminal differentiation. Concurrently, the *Cd68*^+^ SC, *Pdgfra*^+^ SC and *Pecam1*^+^ SC peaked at 1 dpi before modestly declining at 3 dpi, reflecting their transient emergence during the early acute phase of muscle regeneration.

To further model the developmental progression and continuous cell state transitions during early regeneration, we performed pseudotime trajectory analysis. Mapping the cells onto the pseudotime space revealed a bifurcated trajectory, progressing from Early state (predominantly 0/1 dpi cells) toward two distinct fate branches: Late 1 and Late 2, which were primarily populated by cells from the 3 dpi and 1/3 dpi, respectively (Fig. 1E-F). The Early-Late 1 axis accurately captured the canonical myogenic program, showing a significant increase in the proportion of PSC, CSC and DSC, accompanied by a corresponding decrease in QSC and ASC along the pseudotime. Conversely, while the Early-Late 2 branch also exhibited a decline in QSC and ASC, it was distinguished by a significant accumulation of the *Cd68^+^* SC, *Pdgfra^+^* SC and *Pecam1^+^* SC along the trajectory (Fig. 1G and Suppl. Fig. 1B). To delineate the molecular drivers of these divergent fates, we analyzed differentially expressed genes (DEGs) between the Late 1 and Late 2 branches. Hierarchical clustering of the top 20 DEGs revealed distinct transcriptional signatures defining each branch (Fig. 1H and Suppl. Dataset 1). The Late 1 branch was uniquely characterized by the expression of genes governing cell proliferation (*Plk1*, *Kif18b*, *Anln*, *Pimreg*), MuSC differentiation (*Myog*, *Mymk*, *Mymx*, *Cavin4*, *Synpo2l*, *Tnnt2*) and neuromuscular junction assembly (*Chrng*, *Chrnd*, *Dpp6*, *Frmpd1*). In contrast, the Late 2 branch exhibited an enrichment of anti-apoptotic genes (*Bcl2a1a*, *Bcl2a1b*, *Bcl2a1d*), potent inflammatory mediators (*Ccl3*, *Ccl4*, *Il1a*, *Il1b*, *Il7r*, *Il1f9*, *Nlrp3*, *Ptgs2*) and immune response effectors (*Cybb*, *Lcn2*, *S100a8*, *S100a9*). Altogether, these results underscore the emergence of highly heterogeneous cell states within the activated MuSC pool following injury.

Next, to specifically identify the senescent MuSCs during the early regeneration, we employed our recently developed USS method^12^, which utilizes a robust signature of over 500 senescent genes (Suppl. Dataset 1) and demonstrates superior efficacy in defining senescent cells compared to single-marker approaches. Projecting the USS score onto the pseudotime trajectory and UMAP space revealed a distinct subpopulation of cells with elevated USS scores, which were predominantly localized to the Late 2 branch and mapped closely to the *Cd68^+^*, *Pdgfra^+^*subpopulations (Fig. 1I-J). The cells were then segregated into USS^Hi^ and USS^Lo^ groups based on the scoring (Fig. 1K). Quantification across the regeneration timeline showed a progressive expansion of the USS^Hi^ MuSCs population, increasing from 3.1% at 0 dpi to 6.2% at 1 dpi, and peaking at 8.9% by 3 dpi (Fig. 1L). Consistent with the trajectory analysis, USS^Hi^ cells were predominantly enriched within the Cd68 subpopulation at 1 dpi, and both Cd68 SC and Pdgfra SC at 3 dpi (Suppl. Fig. 1C). Differential gene expression (DEG) analysis at both 1 and 3 dpi between USS^Hi^ and USS^Lo^ MuSCs displayed a striking Gene Ontology (GO) enrichment for pathways associated with inflammation and immune regulation, including “inflammatory response”, “immune response”, and “chemotaxis” (Fig. 1M-N and Suppl. Dataset 1). Consistent with these functional annotations, violin plots confirmed that USS^Hi^ MuSCs expressed significantly higher levels of inflammatory components, such as *Ccl2*, *Ccl3*, *Cxcl2*, and *Il1b* (Fig. 1O). Notably, an assessment of commonly used senescent cell markers revealed no detectable expression of *Cdkn2a* (p16^INK4a^) expression in these cells, aligning with several earlier studies^18,23^. Furthermore, *Cdkn2d* (p19) and *Cdkn1a* (p21) expression levels were not consistently elevated in the USS^Hi^ MuSCs at 1 and 3 dpi, raising caution against relying on *Cdkn2a* (p16^INK4a^) or other single markers for the identification of senescent MuSCs (Fig. 1O). Altogether, these findings indicate that the USS^Hi^ subpopulation represents a distinct, senescent-like MuSC state emerging during early muscle regeneration and is characterized by a robust pro-inflammatory transcriptomic signature.

### Isolation and characterization of USS^Hi^ MuSCs reveals a unique senescent-like state

To functionally characterize USS^Hi^ MuSCs and elucidate their function, we sought a robust surface marker to physically isolate these cells (Fig. 2A). Transcriptomic analysis revealed high expression levels of major histocompatibility complex (MHC) class II antigens (*H2-Aa*, *H2-Ab1*, *H2-Eb1*, *H2-DMa*, and *H2-DMb1*) within the USS^Hi^ population (Fig. 2B). Utilizing an MHCII antibody for FACS sorting indeed yielded a clear separation of MHCII+ from MHCII- MuSCs (Fig. 2C and Suppl. Fig. 2A). The percentage of MHCII+ cells increased progressively from 1 to 3 dpi (Fig. 2D), consistent with the dynamics of USS^Hi^ cell population (Fig. 1L). Furthermore, the isolated cells showed a robust upregulation of SASP cytokines and chemokines identified in the USS^Hi^ MuSCs (Fig. 1O), including *Ccl2*, *Ccl3*, *Ccl4*, *Ccl6*, *Il1b*, and *Cxcl2* at 1 dpi and *Ccl2*, *Cxcl1*, *Cxcl2*, *Tnfrsf1a*, *Mmp14* and *Thbs1* at 3 dpi (Fig. 2E-F). These data thus establish MHCII as a reliable surrogate surface marker for isolating the USS^Hi^ MuSCs subpopulation.

**Figure 2.**
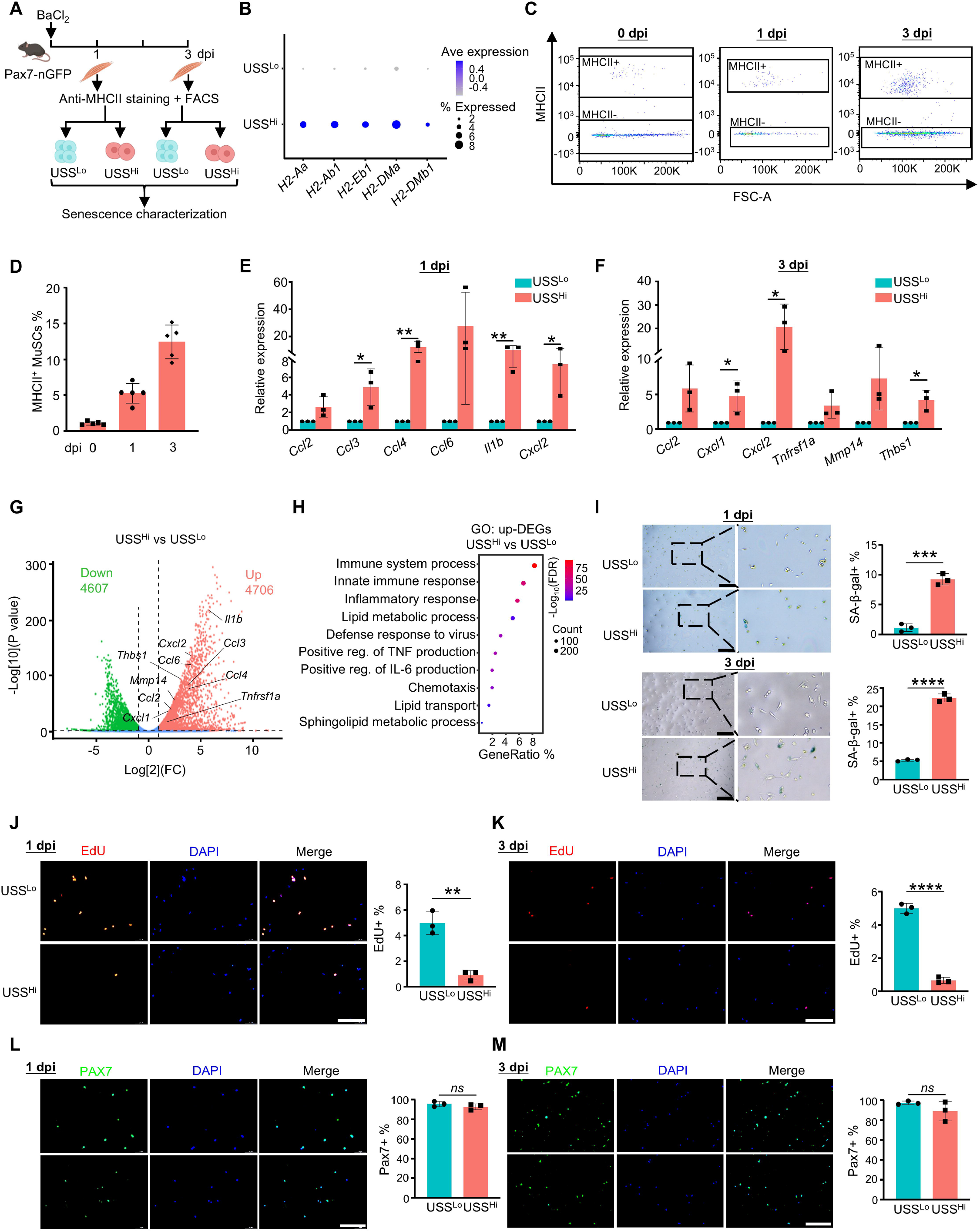
Isolation and characterization of USS^Hi^ MuSCs reveals a unique senescent-like state. **(A)** Schematic of the FACS isolation of USS^Lo^ and USS^Hi^ MuSCs from regenerating muscle at 1 and 3 dpi. **(B)** Dot plot showing the expression of MHC class II genes in USS^Hi^ vs. USS^Lo^ MuSCs. **(C)** Representative FACS plots showing the separation of MHCII+ from MHCII- MuSCs at 0, 1 and 3 dpi. **(D)** Quantification of the MHCII+ MuSC fraction across the regeneration timeline. *n* = 5. **(E-F)** qRT-PCR validation of the selected SASP expression in sorted USS^Hi^ vs. USS^Lo^ MuSCs at 1 dpi (E) and 3 dpi (F). *n* = 3. **(G)** Volcano plot showing the DEGs from bulk RNA-seq performed in the above USS^Hi^ and USS^Lo^ MuSCs. **(H)** GO enrichment analysis of the upregulated DEGs. **(I)** SA-β-gal staining of the above USS^Lo^ and USS^Hi^ MuSCs at 1 and 3 dpi with quantification of the SA-β-gal^+^ cells. Scale bar: 200 μm, *n* = 3. **(J-K)** EdU incorporation assay in the above cells with quantification of EdU+ cells. Scale bar: 150 μm, *n* = 3. **(L-M)** PAX7 immunostaining in the above cells with quantification of PAX7+ cells. Scale bar: 150 μm, *n* = 3. All the bar graphs are presented as mean ± SD, unpaired two-sided Student’s t-test was used to calculate the statistical significance: \**p* <0.05, \*\**p* <0.01, \*\*\**p* <0.001, \*\*\*\**p*<0.0001, *ns* = no significance.

To characterize the above isolated USS^Hi^ cells, we performed bulk RNA-seq to validate their transcriptomic distinction from the USS^Lo^ cells (Suppl. Dataset 2). Consistently, profound gene expression changes were detected in USS^Hi^ vs USS^Lo^ cells, with 4706 genes significantly upregulated and 4607 genes downregulated (Fig. 2G). GO enrichment analysis of the upregulated DEGs reinforced their specialized functional state, highlighting a strong induction of pathways related to “immune system process”, “innate immune response”, and “inflammatory response” (Fig. 2H and Suppl. Dataset 2). Notably, the above SASPs (Fig. 2E-F) were among the highly upregulated genes. Furthermore, SA-β-gal staining revealed a significantly higher percentage of positive cells in the isolated USS^Hi^ MuSCs compared to USS^Lo^ controls at both 1 and 3 dpi (Fig. 2I). Consistently, EdU incorporation assays also showed USS^Hi^ MuSCs exhibited severe impairment in proliferative capacity at both 1 and 3 dpi (Fig. 2J-K), suggesting the USS^Hi^ cells exhibit typical senescent phenotypes. Nevertheless, no significant difference in the PAX7 staining was detected between the USS^Hi^ and USS^Lo^ populations (Fig. 2L-M). Together, these findings indicate that USS^Hi^ MuSCs enter a unique senescent-like state characterized by elevated expression of proinflammatory SASPs, pronounced SA-β-gal activity and diminished proliferation, while maintaining their defining PAX7 identity.

### Multiomics profiling highlights the robust secretory phenotypes and lipid remodeling of USS^Hi^ MuSCs

To further characterize the USS^Hi^ population, we performed multiomics profiling. First, global mass spectrometry (MS)-based proteomic profiling was conducted using whole protein lysates from isolated USS^Lo^ and USS^Hi^ MuSCs at 3 dpi (Fig. 3A and Suppl. Dataset 3). Analysis of the differentially expressed proteins (DEPs) detected a significant intracellular upregulation of key inflammatory SASP mediators in USS^Hi^ cells, including CCL2, IL1b, IL16, TGFb1, MMP14, and TNFRSF1b (Fig. 3B). GO enrichment analysis of the upregulated DEPs (up-DEPs) demonstrated a massive enrichment for biological processes governing immune and inflammatory regulation, such as “regulation of immune response”, “leukocyte activation”, “cytokine production” and “inflammatory response” (Fig. 3C and Suppl. Dataset 3). The above proteomic data tightly parallel our scRNA-seq findings (Fig. 1), confirming that USS^Hi^ MuSCs undergo profound intracellular reprogramming toward a pro-inflammatory state.

**Figure 3.**
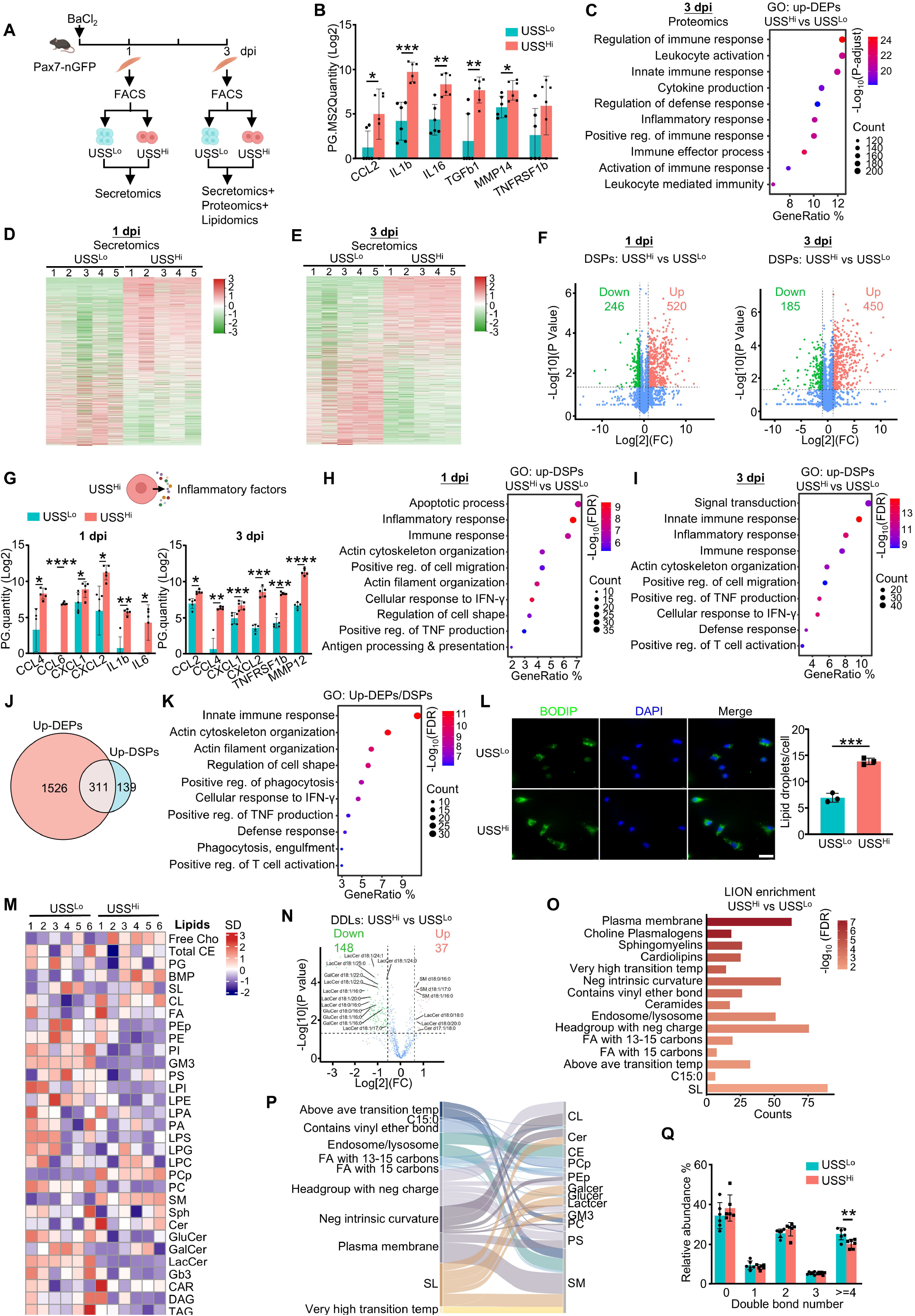
Multiomics profiling highlights the robust secretory phenotypes and lipid remodeling of USS^Hi^ MuSCs. **(A)** Schematic of the multiomics profiling: mass spectrometry (MS)-based proteomics, secretomics and lipidomics were performed on FACS-isolated USS^Lo^ and USS^Hi^ MuSCs at 1 or 3 dpi. **(B)** Intracellular protein abundance (represented by Protein Group MS2 Quantity) from analyzing the proteomics of selected key SASP proteins in USS^Hi^ vs. USS^Lo^ MuSCs at 3 dpi. *n* = 6. **(C)** GO enrichment analysis of the above upregulated DEPs at 3 dpi. **(D-E)** Hierarchical clustering heatmaps defined by the secretome analysis of USS^Lo^ and USS^Hi^ MuSCs at 1 (D) and 3 dpi (E). **(F)** Volcano plots showing the DSPs of USS^Hi^ vs. USS^Lo^ MuSCs at 1 and 3 dpi. **(G)** Secreted abundance (represented by Protein Group Quantity) of selected SASP factors from the above secretomics analysis at 1 and 3 dpi. *n* = 5. **(H-I)** GO enrichment analysis of the above upregulated DSPs at 1 (H) and 3 dpi (I). **(J)** Venn diagram showing the overlap between the upregulated DEPs and DSPs at 3 dpi. **(K)** GO enrichment analysis of the above overlapping proteins. **(L)** BODIPY staining of intracellular lipid droplets in sorted USS^Lo^ and USS^Hi^ MuSCs at 3 dpi, with quantification of lipid droplets per cell. Scale bar: 20 μm, *n* = 3. **(M)** Heatmap of abundances of the indicated lipid classes measured by the lipidomic profiling of USS^Lo^ and USS^Hi^ MuSCs at 3 dpi. *n* = 6. **(N)** Volcano plot showing the differentially detected lipid (DDL) species between USS^Hi^ vs. USS^Lo^ MuSCs. Significantly altered sphingolipid (SL) species are explicitly annotated. **(O)** LION lipid ontology enrichment analysis of the DDLs. **(P)** Sankey diagram linking the biophysical properties of lipids to the DDLs. **(Q)** Relative abundance of lipids stratified by double bond number in USS^Hi^ vs. USS^Lo^ MuSCs. *n* = 6. All the bar graphs are presented as mean ± SD, unpaired two-sided Student’s t-test was used to calculate the statistical significance: \**p* <0.05, \*\**p* <0.01, \*\*\**p* <0.001, \*\*\*\**p*<0.0001, *ns* = no significance.

To demonstrate that USS^Hi^ MuSCs actively secrete the inflammatory protein factors, we performed MS-based secretomic profiling using the conditioned medium harvested from isolated USS^Lo^ and USS^Hi^ MuSCs at both 1 and 3 dpi (Fig. 3A and Suppl. Dataset 4). Principal component analysis (PCA) of the secretomic data revealed a stark separation between the USS^Hi^ and USS^Lo^ subpopulations (Suppl. Fig. 3A). USS^Lo^ MuSCs at 1 dpi and 3 dpi segregated clearly from one another whereas the USS^Hi^ MuSCs at 1 dpi and 3 dpi clustered tightly together, indicating the USS^Hi^ cells maintain a highly stable secretory profile along the regenerative timeline. Hierarchical clustering supported the distinct protein secretion landscapes between the two subpopulations (Fig. 3D-E). Quantitative analysis identified 520 significantly increased and 246 decreased differentially secreted proteins (DSPs) in the USS^Hi^ MuSCs at 1 dpi, and similarly, 450 upregulated and 185 downregulated DSPs at 3 dpi (Fig. 3F). Further examination of these upregulated DSPs (up-DSPs) revealed that USS^Hi^ cells secreted significantly higher quantities of classical inflammatory components, including chemokines (CCL2, CCL4, CCL6, CXCL1, CXCL2), interleukins (IL1b, IL6) and tissue remodeling factors (TNFRSF1b, MMP12) at 1 and 3 dpi (Fig. 3G). Subsequent GO analysis of the up-DSPs consistently highlighted pathways related to “inflammatory response”, “immune response”, and “positive regulation of TNF production”, indicating a sustained and active release of inflammatory proteins from USS^Hi^ cells during early muscle regeneration phase (Fig. 3H-I and Suppl. Dataset 4). Interestingly, GO analysis of the significantly downregulated DSPs (down-DSPs) revealed a marked reduction in proteins associated with “cell adhesion”, “extracellular matrix organization”, and “cell migration” at 1 dpi (Suppl. Fig. 3B). By 3 dpi, the downregulated secretory profile shifted to reflect a suppression of fundamental cellular maintenance, heavily enriched in “positive regulation of transcription”, “actin cytoskeleton organization”, and “protein folding” (Suppl. Fig. 3C).

Next, integrating the 3 dpi proteomic and secretomic datasets (Fig. 3J), we found that 69.1% (311 proteins) of the up-DSPs were also up-DEPs significantly elevated at the intracellular protein level (Suppl. Dataset 5). GO enrichment of these proteins reinforced their core function in immune response related pathway (Fig. 3K and Suppl. Dataset 5).

Finally, we also performed lipidomic profiling (Fig. 3A and Suppl. Dataset 6) reasoning secretory lipids are emerging as important SASP components but remain largely under-investigated. Of note, our earlier transcriptomic GO analysis revealed a significant enrichment of pathways related to “lipid metabolic process”, “lipid transport” and “sphingolipid metabolic process” in the USS^Hi^ subpopulation (Fig. 2H), suggesting a possible alteration in lipid synthesis. Additionally, we observed significant increase of intracellular lipid droplets (LDs) in USS^Hi^ MuSCs compared to their USS^Lo^ counterparts by BODIPY staining of MuSCs sorted at 3 dpi (Fig. 3L), which is consistent with the previous report showing LD accumulation is an established hallmark of cellular senescence^29^. To characterize the specific intracellular lipid species, total lipids were extracted from isolated USS^Lo^ and USS^Hi^ MuSCs at 3 dpi and subjected to MS-based lipidomic analysis. PCA of the resulting dataset revealed a robust separation between the two subpopulations, indicating a fundamental metabolic divergence (Suppl. Fig. 3D). This distinct segregation was visually corroborated by hierarchical clustering, which displayed highly contrasting global lipidomic signatures between the USS^Hi^ and USS^Lo^ cells (Fig. 3M). Quantitative volcano plot analysis detailed the profound lipidomic shift with differentially detected lipid (DDL) species, identifying 148 significantly decreased and 37 increased DDL species in USS^Hi^ MuSCs (Fig. 3N). Notably, a substantial proportion of these top altered species belonged to the sphingolipid (SL) family including Lactosylceramide (LacCer), Glucosylceramide (GluCer), Ceramide (Cer) and Sphingomyelin (SM).

To gain functional insights into this lipidomic remodeling, Lipid Ontology (LION) enrichment analysis^30^ was conducted, which identified significantly enriched functional terms and their corresponding lipid classes in USS^Hi^ MuSCs (Fig. 3O-P). Consistent with our earlier transcriptomic observations and quantitative analysis, the LION analysis prominently highlighted terms related to the SL family, encompassing “Sphingomyelins”, “Ceramides”, and the “Plasma membrane” localization (Fig. 3O). To specifically evaluate this targeted remodeling, we assessed the Log2 fold changes across all detected lipid classes (Suppl. Fig. 3E), which revealed a selective rewiring of SL metabolism: glycosphingolipids, including LacCer and GluCer, were markedly decreased, whereas Cer itself and SM were notably elevated in the USS^Hi^ MuSCs, suggesting that SL flux is redirected away from glycosphingolipid synthesis toward Cer/SM accumulation. Beyond these compositional alterations, the LION analysis intriguingly enriched for terms describing specific biophysical lipid properties, including “Very high transition temperature” and “Above average transition temperature”, alongside the accumulation of fully saturated fatty acids such as “C15:0” (Fig. 3O and Suppl. Dataset 6). The combination of elevated physical transition temperatures and increased lipid saturation indicates a pronounced reduction in overall membrane fluidity and increased membrane stiffness within USS^Hi^ MuSCs. A decline in membrane fluidity is a critical biophysical feature of cellular senescence, often driving the formation of rigid lipid microdomains that fundamentally alter cell signaling and receptor dynamics^31,32^. This biophysical adaptation is further corroborated by the quantitative assessment of lipid saturation (Fig. 3Q), which demonstrated a significant decrease in the relative abundance of highly polyunsaturated lipids (≥4 double bonds) in the USS^Hi^ MuSCs, a structural remodeling that functionally restricts membrane fluidity.

Collectively, this multi-omics profiling uncovers that the USS^Hi^ MuSC subpopulation acquires a potent pro-inflammatory protein secretory phenotype coupled with sphingolipid remodeling during early skeletal muscle regeneration.

### Targeted *in vivo* clearance of USS^Hi^ MuSCs impairs skeletal muscle regeneration and stem cell expansion

To pinpoint the precise function of USS^Hi^ MuSCs *in vivo* and address conflicting views regarding cellular senescence in muscle regeneration, a genetic approach specifically removing this cell population is essential. Since commonly used p16^INK4a^-based ablation strategies were inapplicable due to a lack of *Cdkn2a* (p16^INK4a^) expression (Fig. 1O), we developed a genetic ablation strategy based on the observation that *Ccl2* is highly induced in USS^Hi^ MuSCs but barely detectable in USS^Lo^ cells (Fig. 1O); moreover, USS^Hi^ MuSCs exhibit elevated production and secretion of CCL2 protein (Fig. 3B, G). We knocked in the Loxp-stop codon-Loxp-diphtheria toxin A (DTA) after the exon 3 of *Ccl2* gene and crossed this *Ccl2-LSL-DTA* mice with *Pax7-CreERT* and *Rosa26-LSL-YFP* reporter mouse strain^27^ to generate a triple-transgenic Knock-In (KI) model (Fig. 4A). Upon tamoxifen (TMX) administration, DTA is selectively expressed in cells concurrently expressing *Pax7* and *Ccl2*, thereby inducing specific clearance of the USS^Hi^ MuSCs from the early stage of regeneration.

**Figure 4.**
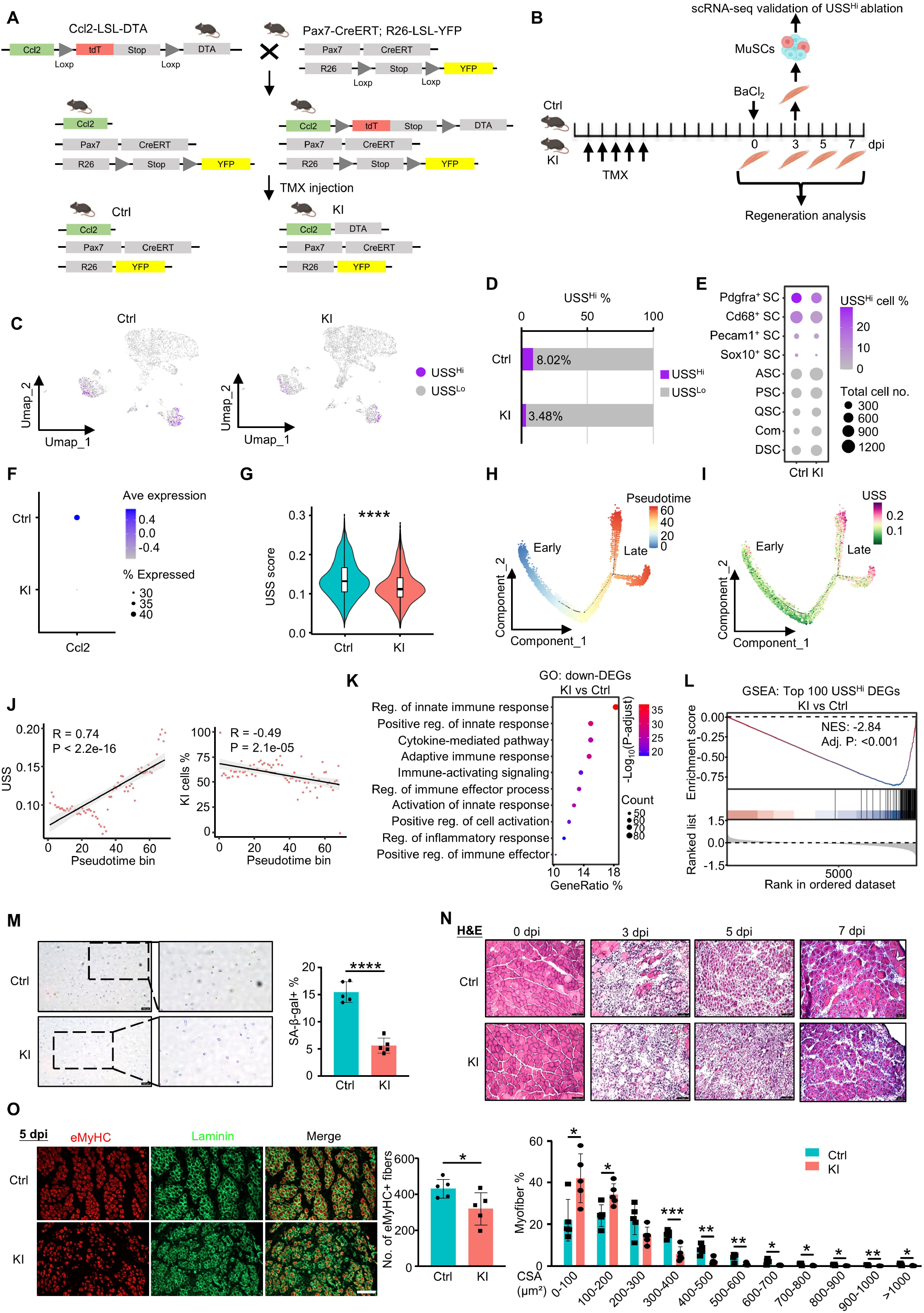
Targeted *in vivo* clearance of USS^Hi^ MuSCs impairs skeletal muscle regeneration and stem cell expansion. **(A)** Schematic representation of the *Ccl2*-DTA knockin (KI) mediated strategy used to generate the USS^Hi^ MuSC ablation mouse model. **(B)** Experimental timeline for ablation validation in the KI mice and subsequent regeneration analysis. **(C)** UMAP plots highlighting the distribution and density of USS^Hi^ and USS^Lo^ populations in the Ctrl (left) and KI (right) groups. **(D)** Percentage of USS^Hi^ cells in Ctrl vs. KI mice. **(E)** Dot plot illustrating the percentage of USS^Hi^ MuSCs and total cell number distributed across the various annotated MuSC clusters in both Ctrl and KI conditions. **(F)** *Ccl2* expression in MuSCs of Ctrl and KI mice. **(G)** USS score distribution in MuSCs of Ctrl and KI mice. **(H-I)** Pseudotime trajectory of MuSCs colored by pseudotime (H) and by USS (I). **(J)** Scatter plots showing the Spearman correlation between USS score and KI cells percentage along the pseudotime bins. **(K)** GO enrichment analysis of downregulated DEGs in KI vs. Ctrl MuSCs. **(L)** GSEA of the top 100 USS^Hi^ DEGs in KI versus Ctrl MuSCs. **(M)** SA-β-gal staining of sorted MuSCs from Ctrl and KI mice at 3 dpi, with quantification. Scale bar: 100 μm, *n* = 5. **(N)** H&E staining of TA muscle sections at 0, 3, 5 and 7 dpi in Ctrl and KI mice. Scale bar: 75 μm, *n* = 5. **(O)** Immunofluorescence staining of eMyHC and Laminin at 5 dpi, with quantification of eMyHC^+^ fiber number and cross-sectional area (CSA) distribution. Scale bar: 100 μm, *n* = 5. All the bar graphs are presented as mean ± SD, unpaired two-sided Student’s t-test was used to calculate the statistical significance: \**p* <0.05, \*\**p* <0.01, \*\*\**p* <0.001, \*\*\*\**p*<0.0001, *ns* = no significance.

To validate the efficiency of this ablation strategy, MuSCs were isolated from Control (Ctrl) and KI mice at 3 dpi and subjected to scRNA-seq (Fig. 4B, Suppl. Fig. 4A-C and Suppl. Dataset 7). UMAP projections and cell cluster quantification confirmed a successful and dramatic depletion of the USS^Hi^ population, which dropped from 8.02% in Ctrl to 3.48% in KI mice (Fig. 4C-D). Consistent with the initial scRNA-seq analysis (Fig. 1G, I-K and Suppl. Fig. 1C), USS^Hi^ MuSCs in control mice were predominantly enriched within the *Pdgfra*^+^ SC and *Cd68*^+^ SC, and following the DTA-mediated ablation in KI mice, the proportion of USS^Hi^ cells within these *Pdgfra*^+^ SC and *Cd68*^+^ SC exhibited a striking reduction but other canonical MuSC subclusters, QSC, ASC, PSC, CSC and DSC, remained largely unperturbed (Fig. 4E and Suppl. Fig. 4B). Concomitantly, the overall expression of *Ccl2* (Fig. 4F) and the global USS score (Fig. 4G) were significantly diminished in the KI group. Subsequent pseudotime trajectory analysis revealed a bifurcated trajectory containing two “Late” branches, with the USS^Hi^ MuSCs exclusively mapping to the distal termini of the late branches (Fig. 4H-I). Notably, a robust positive correlation between pseudotime progression and the cellular USS was identified (*R*=0.74, *P*<2.2e-16) but the fraction of MuSCs derived from KI mice exhibited a significant negative correlation with pseudotime (*R*=-0.49, *P*=2.1e-05) (Fig. 4J). This inverse correlation confirms the targeted elimination of USS^Hi^ MuSCs in the KI mouse model. GO enrichment analysis of the significantly downregulated genes in the KI MuSCs demonstrated the abrogation of pathways governing the pro-inflammatory secretory phenotype of USS^Hi^ MuSCs (Fig. 4K and Suppl. Dataset 7). Moreover, Gene Set Enrichment Analysis (GSEA) utilizing the top 100 defining signature genes of the USS^Hi^ MuSCs (Suppl. Dataset 7) yielded a significant negative enrichment within the KI dataset (NES =-2.84, Adj. *P* < 0.001) (Fig. 4L), confirming the suppression of the USS^Hi^ transcriptional signature in KI mice. To further validate the clearance of USS^Hi^ MuSCs, *in vitro* SA-β-gal staining of sorted MuSCs at 3 dpi revealed a significant reduction in the proportion of SA-β-gal^+^ cells in KI vs Ctrl mice (Fig. 4M). Together, these data demonstrate the successful and specific *in vivo* ablation of USS^Hi^ MuSCs in the KI mice.

We next assessed the impact of USS^Hi^ MuSC depletion on the overall kinetics of muscle repair following BaCl_2_ injury (Fig. 4B). Histological evaluation via Hematoxylin and Eosin (H&E) staining at 0, 3, 5, and 7 dpi revealed a marked delay in the regenerative process in KI mice (Fig. 4N): while Ctrl muscles exhibited robust formation of newly regenerating fibers characterized by centrally localized nuclei (CLN) by 5 dpi and extensive architectural restoration by 7 dpi, KI muscles displayed persistent necrotic fibers, excessive interstitial cellular infiltration, and noticeably smaller newly formed myofibers. To further assess this regenerative deficit, muscle sections were immunostained at 5 dpi for embryonic Myosin Heavy Chain (eMyHC), a marker of newly formed myofibers, alongside Laminin (Fig. 4O), which uncovered a significant decreased number of eMyHC^+^ fibers and leftward shift in the cross-sectional area (CSA) distribution of eMyHC^+^ fibers in KI mice, indicating that myofiber growth and maturation were blunted in the absence of the USS^Hi^ subpopulation.

Collectively, these findings indicate that USS^Hi^ MuSCs are not a detrimental pathological byproduct, but rather an indispensable physiological player required for successful skeletal muscle regeneration after acute injuries.

### High-resolution spatial and single-cell transcriptomics reveal that USS^Hi^ MuSCs orchestrate the regenerative microenvironment

Given the robust secretory phenotype of USS^Hi^ MuSCs, we speculated that they promote regeneration by actively modulating the local regenerative microenvironment. To test the notion, we applied Visium HD spatial transcriptomics to Tibialis Anterior (TA) muscles from Ctrl and KI mice at 3 dpi (Suppl. Dataset 8). A dual-resolution analysis approach utilizing both 8 µm bin resolution and single-nuclei segmentation was leveraged to map cellular states directly onto the tissue architecture (Fig. 5A). Guided by classical pathological features typically observed in muscle H&E staining, we delineated the tissue section into three distinct functional zones (Fig. 5B): (1) the “Injury” core, defined by massive interstitial cell infiltration and a near-complete absence of intact muscle fibers; (2) the “Peri-injury” zone, characterized by the concurrent presence of infiltrating interstitial cells and surviving muscle fibers; and (3) the “Non-injury” zone, exhibiting intact muscle fibers with minimal cellular infiltration. Spatial UMAP projections on 8 µm bin resolution revealed that these three pathologically defined zones segregated into starkly distinct transcriptomic clusters (Fig. 5C), validating the spatial zonation strategy to investigate molecular microenvironments.

**Figure 5.**
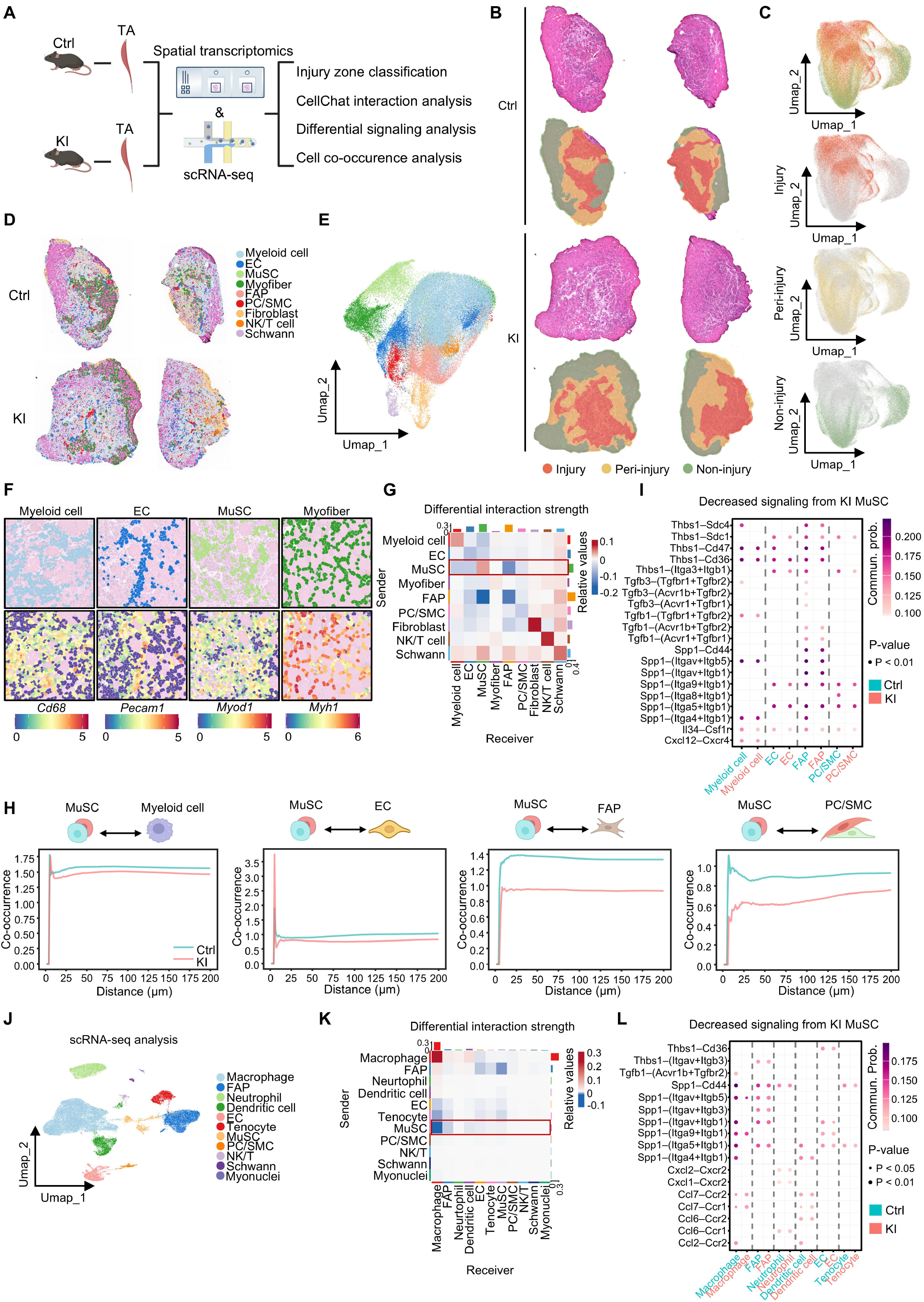
High-resolution spatial and single-cell transcriptomics reveal that USS^Hi^ MuSCs orchestrate the regenerative microenvironment. **(A)** Schematic of the Visium HD spatial transcriptomics and single-cell transcriptomics applied to TA muscles from Ctrl and KI mice at 3 dpi. **(B)** H&E-guided spatial delineation of the “Injury”, “Peri-injury” and “Non-injury” zones in the above Ctrl and KI muscles. **(C)** Spatial UMAP projections of the 8 µm bins resolution spatial transcriptomic data. The top plot displays all spatial 8 µm bins integrated and color-coded by their annotated tissue zones. The subsequent three UMAP plots individually highlight the distribution of transcriptomic profiles corresponding to the “Injury”, “Peri-injury”, and “Non-injury” domains. **(D)** Spatial distribution maps of annotated segmented single nuclei mapped onto the muscle cross-sections from Ctrl and KI mice. **(E)** UMAP embedding of the integrated segmented single nuclei colored by annotated cell type. **(F)** High-resolution spatial mapping validating the cell type annotations. The top row displays the specific spatial localization of individual cell populations on representative tissue sections. The bottom row presents spatial feature plots showing the expression levels of corresponding representative marker genes for each cell lineage. **(G)** CellChat differential interaction strength analysis between Ctrl and KI, showing sender-receiver communications. MuSC emanated signalings to niche cell populations are highlighted. **(H)** Spatial co-occurrence line plots evaluating the co-occurrence probability between MuSCs and Myeloid cells, ECs, FAPs, or Pericytes/SMC across continuous spatial distances. **(I)** Bubble plot of ligand-receptor pairs showing decreased signalings from MuSCs to the designated receiver cells in KI vs. Ctrl. **(J)** UMAP projection of an independent scRNA-seq dataset from Ctrl and KI mice at 1 and 3 dpi. **(K)** CellChat differential interaction strength analysis between Ctrl and KI single-cell transcriptomics data at 3 dpi. MuSC emanated signalings to niche cell populations are highlighted. **(L)** Bubble plot of ligand-receptor pairs showing decreased signalings from MuSCs to the designated receiver cells in KI vs. Ctrl single-cell transcriptomics data at 3 dpi. Bubble plot of decreased signaling from KI MuSCs in scRNA-seq, including chemokine interactions such as *Cxcl1/2-Cxcr2* and *Ccl2/7-Ccr2*.

We next restricted our downstream analysis specifically to the actively regenerating “Injury” area. GO enrichment analysis performed on the 8 µm bin resolution dataset revealed the upregulated genes in the KI muscle were highly enriched for inflammatory and immune-related terms, such as “regulation of innate immune response”, “regulation of immune effector process”, and “regulation of inflammatory response” (Suppl. Fig. 5A and Suppl. Dataset 8). Conversely, the significantly downregulated genes were overwhelmingly enriched for pathways essential to muscle repair and structural maturation, prominently including “muscle system process”, “muscle cell differentiation”, and “striated muscle cell differentiation” (Suppl. Fig. 5A). Consistent with the GO signatures, spatial mapping revealed that essential myogenic regulatory factors (*Myod1*, *Myog*) and the *de novo* regenerating myofiber marker *Myh3* were robustly induced in the Ctrl injury zone but nearly eradicated in the KI muscle (Suppl. Fig. 5B-C). In contrast, the pan-leukocyte marker *Ptprc* exhibited a persistent and robust spatial accumulation within the KI injury zone (Suppl. Fig. 5B-C). This transcriptomic signature implies that the absence of USS^Hi^ MuSCs retards the essential microenvironmental transition from the initial inflammatory phase to the reparative myogenic phase.

Subsequently, utilizing the segmented cell resolution, we mapped the precise spatial distribution of major cell types according to their marker genes, including Myeloid cell, EC, MuSC, Myofiber, and FAP (Fig. 5D-E and Suppl. Fig. 5D). The fidelity of this single-cell spatial annotation was also confirmed by the highly specific, localized expression of canonical markers such as *Cd68*, *Pecam1*, *Myod1*, and *Myh1* within their respective cell lineages (Fig. 5F and Suppl. Fig. 5E-F). The overall intercellular communication landscapes were then evaluated utilizing the CellChat v2 analytical framework. The differential interaction strength network revealed the strength of communication emanating from MuSC to the critical supportive niche cell populations, including Myeloid cell, EC, FAP, and PC/SMC (Pericytes/smooth muscle cells), was substantially diminished upon USS^Hi^ ablation (Fig. 5G and Suppl. Dataset 8), confirming the hypothetical role of USS^Hi^ cells in mediating the MuSC-niche cell crosstalk to modulate the regeneration. To substantiate the above notion, we calculated the spatial co-occurrence probability to assess the physical proximity between MuSCs and these receiver cells. Expectedly, MuSC in KI mice exhibited a markedly reduced physical colocalization thus lower possibility of cross-talking with Myeloid cell, EC, FAP, and PC/SMC measured at various distances (Fig. 5H and Suppl. Dataset 8). Next, in-depth analysis into the specific ligand-receptor interactions responsible for the decreased communication was further conducted and revealed that signaling from MuSC to the surrounding receiving stroma and immune cells via key matrix-remodeling and immunomodulatory pathways was significantly abrogated, including pathways mediated by *Thbs1*, *Tgfb1*, *Tgfb3*, *Spp1*, *Il34* and *Cxcl12* (Fig. 5I). The disruption of *Thbs1* and *Spp1* signaling is known to impact EC and PC/SMC through their CD47 receptors and diverse integrin complexes^33,34^, pointing to a possible defect in angiogenesis and vascular stabilization in KI muscle. Concurrently, the repression of *Tgfb1/3* and *Spp1* to receiving FAP may deprive the microenvironment of vital cues required for proper extracellular matrix (ECM) remodeling and structural scaffolding^35,36^. Given that these specific ligands are also critical drivers of anti-inflammatory M2 macrophage polarization^37–40^, their reduction also explains why the loss of USS^Hi^ MuSCs delayed the essential transition from an unresolved inflammatory state to the reparative myogenic phase (Suppl. Fig. 5A-C). Altogether these data demonstrate that USS^Hi^ ablation impairs MuSCs interactions with multiple niche cells via various paracrine signaling.

To cross-validate our spatial findings and achieve a deeper transcriptomic coverage of these transient signaling events, we performed analogous CellChat analysis on an independently obtained scRNA-seq dataset comparing Ctrl and KI mice at 1 and 3 dpi (Fig. 5J, Suppl. Fig. 5G-H and Suppl. Dataset 9). In concordance with the spatial transcriptomics data, the differential interaction heatmap analysis confirmed a profound, systemic decrease in signaling strength originating from KI MuSCs towards various niche receiver cells at 3 dpi, most notably Macrophages, FAPs, Dendritic cells, and ECs, despite the absence of dramatic changes at 1 dpi (Fig. 5K, Suppl. Fig. 5I and Suppl. Dataset 9). Furthermore, specific evaluation of the impaired signaling pathways at single-cell resolution accurately mirrored the spatial results (*Thbs1*, *Spp1*), while further uncovering a dramatic loss of classical chemokine interactions (*Cxcl1/2-Cxcr2*, *Ccl2/7-Ccr2*) mediated by MuSC (Fig. 5L).

Collectively, these congruent multi-omics analyses provide compelling evidence that USS^Hi^ MuSCs function as crucial signaling hubs via their localized secretory phenotypes, actively cross-talking to multiple niche cells including Myeloid cell, EC, FAP and PC/SMC to orchestrate effective regeneration progression.

### S1P is a key lipid factor mediating USS^Hi^-niche cell communication

To further uncover the key factor(s) mediating the niche-modulatory function of USS^Hi^ cells, we turned our focus to the identified lipid factors (Fig. 3M-O) reasoning the essential function of lipid SASP factors in mediating senescent cell function is under-investigated compared to the protein SASP factors. To pinpoint the potential key SL family lipids, we integrated our lipidomic data with matched transcriptomic and proteomic profiles (Fig. 2G and Fig. 3B) and mapped these datasets onto the canonical SL metabolic pathway. A highly coordinated molecular signature was observed showing that the catabolism of complex SLs into bioactive end-products, Sphingosine-1-phosphate (S1P), was highly elevated in USS^Hi^ vs USS^Lo^ cells (Fig. 6A). Specifically, the enzymes responsible for the sequential degradation of LacCer and GluCer into Ceramide (*Glb1*, *Gba1*, *Gba2*) and the subsequent conversion of Ceramide to Sphingosine (*Asah1*, *Asah2*, *Acer2*) displayed dynamic upregulations at both mRNA and protein levels in USS^Hi^ cells. Concomitantly, the terminal kinases *Sphk1* and *Sphk2*, which sphingosine to produce S1P, were consistently upregulated across transcriptomic and proteomic analyses. This unified multi-omics landscape indicates a unidirectional metabolic flux driving the aberrant synthesis and accumulation of S1P in USS^Hi^ MuSCs. S1P is a potent bioactive signaling lipid that can be exported to the extracellular space to mediate diverse biological responses. Given the metabolic rewiring observed, we hypothesized that USS^Hi^ cells might actively secrete S1P as a lipid-based component of their secretory phenotype. Indeed, S1P enzyme-linked immunosorbent assay (ELISA) of the conditioned medium confirmed that USS^Hi^ MuSCs secreted significantly higher concentrations of S1P compared to USS^Lo^ cells (Fig. 6B). Collectively, the above results demonstrate that the heightened production and secretion of bioactive lipid S1P may mediate the niche-modulating function of USS^Hi^ MuSCs in muscle regeneration.

**Figure 6.**
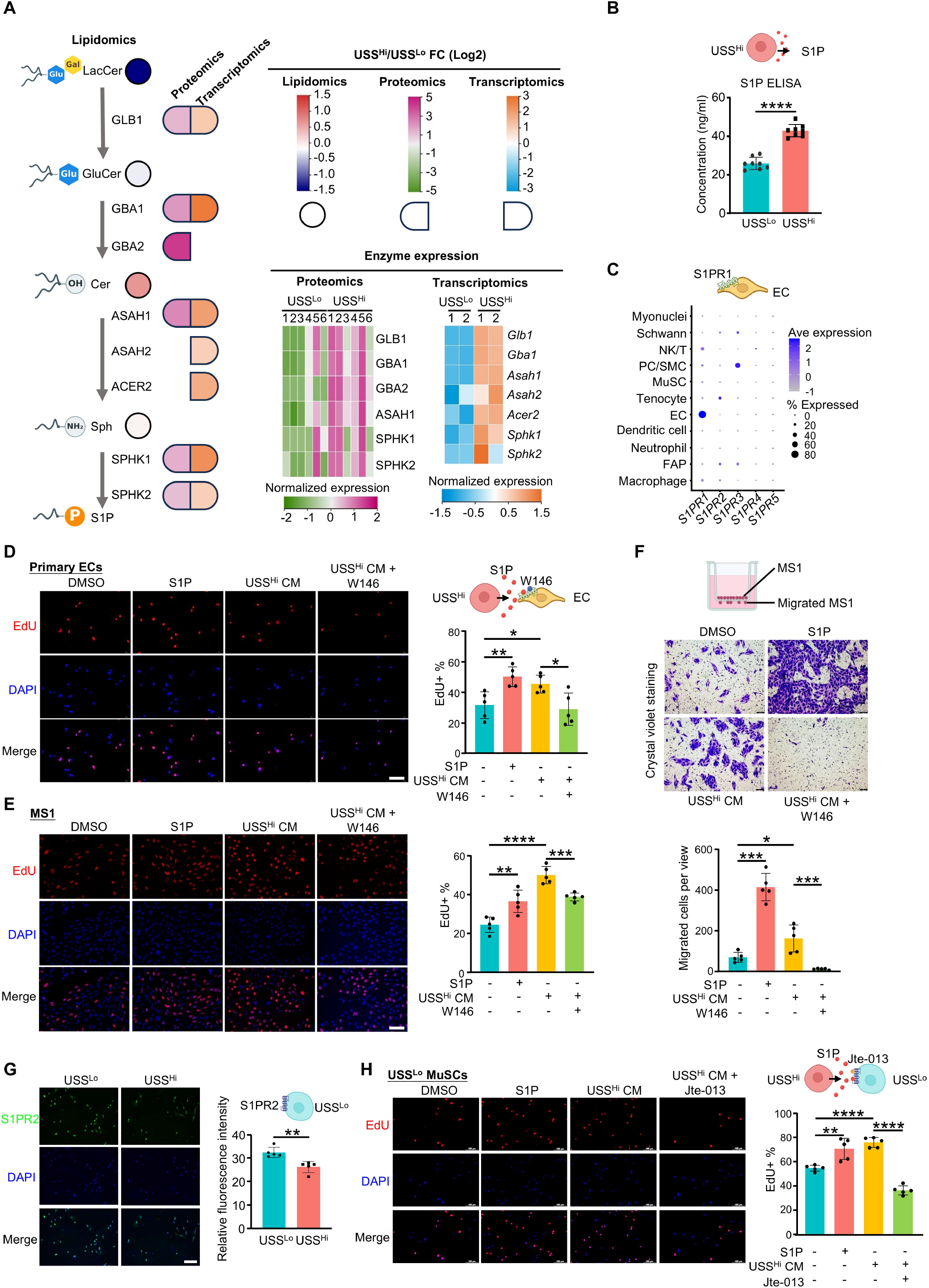
S1P is key lipid factor mediating USS^Hi^-niche cell communication. **(A)** Integrated multiomics analysis of glycosphingolipid catabolic pathway from LacCer to S1P, combining lipidomic, proteomic, and transcriptomic data. Circles represent lipid metabolites, color-coded by their Log2(fold change) (USS^Hi^/USS^Lo^) from the lipidomic dataset, where blue color indicates a decrease and red indicates an increase in USS^Hi^ MuSCs. The split capsule symbols adjacent to the enzymatic steps represent the Log2(fold change) of the corresponding enzymes derived from proteomics (left half; green to pink scale) and transcriptomics (right half; blue to orange scale). The heatmaps on the right display the normalized relative expression levels of key metabolic enzymes (GLB1, GBA1, GBA2, ASAH1, ASAH2, ACER2, SPHK1, and SPHK2) across individual biological replicates in the proteomic and transcriptomic datasets. **(B)** ELISA detected concentration of S1P in the conditioned medium collected from USS^Lo^ and USS^Hi^ MuSCs. **(C)** Dot plot of S1P receptor (*S1pr1-5*) expression across the indicated cell types of the regenerating muscle from analyzing the scRNA-seq data, highlighting the EC-specific expression of S1pr1. **(D)** Primary ECs were isolated from TA muscle at 3 dpi and treated with DMSO, S1P, USS^Hi^ conditioned medium, or USS^Hi^ conditioned medium plus the S1PR1 antagonist W146. EdU assay was performed with quantification of the EdU^+^ cells. Scale bar: 100 μm, *n* = 5. **(E)** EdU staining assay was performed on MS1 cells under the above treatments with quantification of the EdU^+^ cells. Scale bar: 100 μm, *n* = 5. **(F)** Transwell migration assay was performed on MS1 endothelial cell line under the above treatments, and crystal violet staining was performed with quantification of migrated cells per view (lower panel). Scale bar: 100 μm, *n* = 5. **(G)** Immunofluorescence staining of S1PR2 in USS^Lo^ and USS^Hi^ MuSCs at 3 dpi, with quantification of relative fluorescence intensity. Scale bar: 100 μm, *n* = 5. **(H)** USS^Lo^ MuSCs were treated with DMSO, S1P, USS^Hi^ conditioned medium, or USS^Hi^ conditioned medium plus the S1PR2 antagonist JTE-013 and EdU assay was performed with quantification of the EdU^+^ cells Scale bar: 100 μm, *n* = 5. All the bar graphs are presented as mean ± SD, unpaired two-sided Student’s t-test was used to calculate the statistical significance: \**p* <0.05, \*\**p* <0.01, \*\*\**p* <0.001, \*\*\*\**p*<0.0001, *ns* = no significance.

Knowing that S1P is a crucial lipid mediator that maintains EC integrity, promotes vascular maturation, and strengthens the endothelial barrier^41^, we suspected that S1P may play a key role in mediating the identified USS^Hi^-EC cross-talk (Fig. 5G). Indeed, examining the expression of S1P receptors (*S1pr1-5*) by the scRNA-seq analysis revealed *S1pr1* is robustly and specifically expressed on ECs of the regenerating muscle (Fig. 6C), suggesting a targeted paracrine communication axis from USS^Hi^ MuSCs to the local vasculature through S1P-S1PR1. To validate this hypothesis, we isolated primary ECs from regenerating muscles at 3 dpi (Suppl. Fig. 6A) and found both S1P treatment and USS^Hi^ conditioned medium (CM) robustly promoted the proliferation of these primary ECs, a phenotype that was effectively neutralized by the S1PR1-specific antagonist W146 (Fig. 6D). Parallel validation experiments utilizing the mouse EC cell line MS1 were also performed, which confirmed that S1P signaling via S1PR1 not only drove EC proliferation (Fig. 6E), but also significantly enhanced EC migration assessed by Transwell assays (Fig. 6F).

Of note, previous studies showed that S1P acts directly on MuSCs via S1PR2 to promote myoblast proliferation and regeneration^42^. We thus speculated that S1P may also facilitate USS^Hi^-USS^Lo^ cell communication and myoblast expansion. Consistent with this idea, IF staining demonstrated that a higher expression of S1PR2 in USS^Lo^ vs. USS^Hi^ MuSCs (Fig. 6G), indicating a potential regulatory influence of USS^Hi^ MuSCs on USS^Lo^ MuSCs via S1P-S1PR2. Furthermore, treatment of USS^Lo^ MuSCs with either exogenous S1P or USS^Hi^ CM significantly enhanced the cell proliferation, which was completely abrogated by the addition of JTE-013, a specific antagonist for the S1PR2 receptor (Fig. 6H).

Altogether, these findings uncover USS^Hi^ MuSCs reprogram the SL metabolism to secrete high amounts of S1P that facilitates paracrine interaction with ECs and USS^Lo^ MuSCs in the regenerative niche to promote both EC and MuSC expansion.

### S1P is important for USS^Hi^-niche cell communication and muscle regeneration

To further verify the above defined USS^Hi^-niche cell interactions *in vivo*, we first examined the impact of USS^Hi^ depletion on EC and MuSC expansion during muscle regeneration (Fig. 7A). Flow cytometric isolation revealed a significant decrease in the percentage of CD45^-^ & CD31^+^ ECs in KI compared to Ctrl muscles (Fig. 7B). IF staining for CD31 confirmed the marked reduction of EC abundance and vascular deficit of KI mice at 3 dpi (Fig. 7C). Similarly, to validate the USS^Hi^ function on USS^Lo^ expansion *in vivo*, we quantified the MuSC pool at various regenerative stages using flow cytometry. Notably, baseline MuSC numbers at 0 dpi and the initial activation at 1 dpi showed no significant differences between Ctrl and KI mice (Suppl. Fig. 7A), suggesting that the initial response to injury was intact. However, the requisite expansion of the MuSC pool by 3 dpi was significantly compromised in KI mice, yielding a significantly lower percentage of MuSCs compared to Ctrl counterparts (Fig. 7D). Consistently, *in situ* IF staining for PAX7 and MYOD at 3 dpi muscle sections also revealed a significantly reduced number of PAX7^+^ (Fig. 7E) and MYOD^+^ cells (Fig. 7F) in the KI muscle sections. Altogether, the above results demonstrate that USS^Hi^ MuSCs play a critical role in promoting local angiogenesis and MuSC expansion in muscle regeneration.

**Figure 7.**
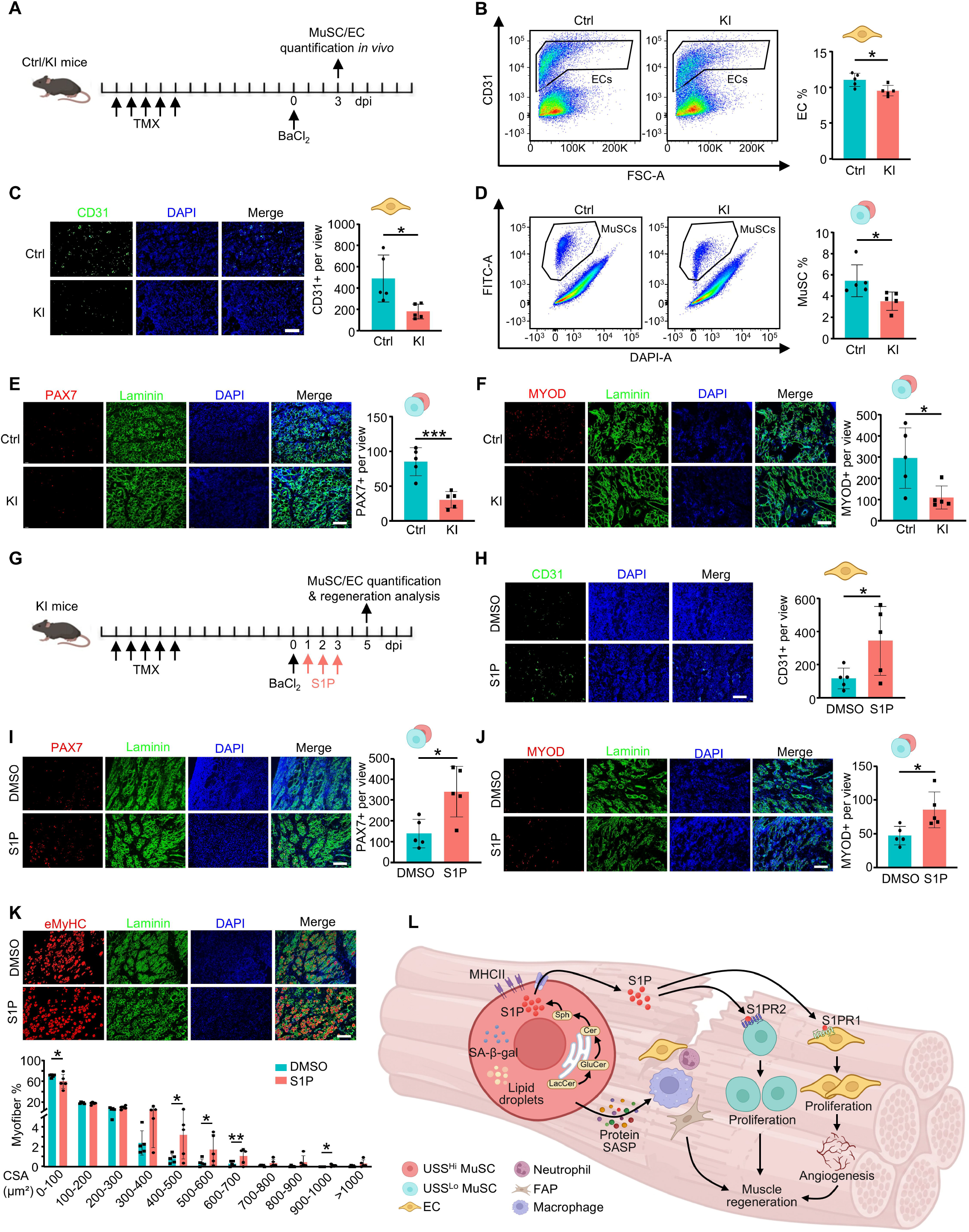
S1P is important for USS^Hi^-niche cell communication and muscle regeneration. **(A)** Schematic of the experimental timeline in Ctrl and KI mice. **(B)** Flow cytometric quantification of CD45- CD31+ ECs in Ctrl and KI muscles at 3 dpi. **(C)** CD31 immunofluorescence staining of Ctrl and KI muscle sections at 3 dpi, with quantification of CD31+ numbers per view. Scale bar: 100 μm, *n* = 5. **(D)** Flow cytometric quantification of the MuSC percentage in Ctrl and KI muscles at 3 dpi. **(E-F)** Immunofluorescence staining of PAX7/Laminin (E) and MYOD/Laminin (F) in Ctrl and KI muscle sections at 3 dpi with quantification of PAX7+ and MYOD+ cells per view. Scale bar: 100 μm, *n* = 5. **(G)** Schematic of the *in vivo* S1P rescue experiment in KI mice following BaCl_2_ injury. **(H)** CD31 immunofluorescence staining of S1P- vs. DMSO-treated KI muscles, with quantification of CD31+ numbers per view. Scale bar: 100 μm, *n* = 5. **(I-J)** PAX7/Laminin (I) and MYOD/Laminin (J) immunofluorescence staining of the above muscles, with quantification of PAX7+ and MYOD+ cells per view. Scale bar: 100 μm, *n* = 5. **(K)** eMyHC/Laminin immunofluorescence staining of the above muscles at 5 dpi, with quantification of the CSA distribution of eMyHC+ regenerating myofibers. Scale bar: 100 μm, *n* = 5. **(L)** Graphical model summarizing the study: senescence-like USS^Hi^ MuSCs drive muscle regeneration through secretory niche orchestration. All the bar graphs are presented as mean ± SD, unpaired two-sided Student’s t-test was used to calculate the statistical significance: \**p* <0.05, \*\**p* <0.01, \*\*\**p* <0.001, \*\*\*\**p*<0.0001, *ns* = no significance.

To solidify that S1P functions as a key factor mediating the promoting effect of USS^Hi^ MuSCs in muscle regeneration, we performed an *in vivo* rescue experiment by administering exogenous S1P following muscle injury in KI mice (Fig. 7G). As expected, exogenous S1P replenished the compromised vascular network, as evidenced by the rescued CD31^+^ EC numbers in the S1P-treated group (Fig. 7H). The PAX7^+^ and MYOD^+^ MuSC pool at 5 dpi was also markedly replenished (Fig. 7I-J). As a result, S1P supplementation was sufficient to reverse the compromised regeneration of the KI mice: Histological assessment at 5 dpi revealed a robust restoration of newly formed, eMyHC^+^ regenerating myofiber CSA in S1P-treated KI mice (Fig. 7K). Collectively, these *in vivo* findings provide compelling evidence that S1P is crucial in enabling the critical niche cell communication and microenvironmental orchestration of USS^Hi^ MuSCs in muscle regeneration (Fig. 7L).

## Discussion

Driven by the motivation of resolving the controversial roles of cellular senescence in skeletal muscle regeneration, the present study identifies a unique population of senescence-like MuSCs, USS^Hi^ cells, which emerge in the early phase of regeneration after acute injury in young mice. By leveraging MHCII mediated FACS isolation, high-resolution multi-omics characterization, and CCL2 mediated cell specific genetic ablation, our data reveal that these cells exert a beneficial role in promoting acute injury induced muscle regeneration through their heightened secretory phenotypes. Through an array of protein/lipid secretomics, the cells act as an active signaling hub cross-talking to multiple cell types in the regenerating niche microenvironment. Specifically, the secreted S1P lipid mediates their communication with ECs and USS^Lo^ MuSCs to orchestrate angiogenesis and myoblast proliferation thereby enabling effective regeneration.

### Cdkn2a-independent identification of USS^Hi^ MuSCs and genetic validation of their pro-regenerative function

A fundamental barrier to understanding senescence in highly plastic tissues has been the field’s over-reliance on canonical markers, most notably *Cdkn2a* (p16^INK4a^), to define the senescent state. Our single-cell transcriptomic mapping of the acute regeneration timeline reveals a critical limitation of this classical approach: the USS^Hi^ MuSCs, despite exhibiting pronounced senescence hallmarks such as elevated SA-β-gal activity, profound cell-cycle arrest, and a robust SASP, are virtually devoid of *Cdkn2a* (p16^INK4a^) expression. Furthermore, the expression of *Cdkn1a* (p21) and *Cdkn2d* (p19) proved inconsistent and insufficient for accurate lineage tracing. This critical observation elucidates why generalized senolytic therapies or transgenic ablation models relying strictly on the *Cdkn2a* (p16^INK4a^) promoter indiscriminately eliminate disparate supporting lineages while simultaneously failing to target this specialized MuSC subpopulation. Indeed, emerging studies highlight the heterogeneity of cellular senescence and the employment of a comprehensive scoring approach is becoming necessary for senescent cell identification. By utilizing USS alongside the identification of MHCII as a reliable surface surrogate, we established a framework to interrogate these cells with unprecedented specificity. The subsequent deployment of our *Pax7-CreERT; Ccl2-LSL-DTA* intersectional genetic model provided the first direct *in vivo* demonstration that the targeted ablation of this specific USS^Hi^ MuSC pool derails the myogenic cascade. Although we have shown the USS^Hi^ MuSCs are β-Gal positive and proliferatively arrested resembling typical senescent cells, we are uncertain about the long-term fate of these cells. It remains a possibility that the cell cycle arrest is transient and reversible, which warrants the implementation of longitudinal lineage tracing to dynamically track the fate and ultimate outcome of these USS^Hi^ MuSCs. It is thus plausible to name the population senescent-like cells.

Our finding that senescence-like USS^Hi^ MuSCs actively promote skeletal muscle regeneration appears to exhibit certain discrepancies with a previous study by Moiseeva et al. that has characterized cellular senescence as a predominantly detrimental presence in the muscle regenerative niche in young mice^20^. This divergence in outcomes may be largely attributable to the specific senescent cell populations targeted and the precision of the ablation strategies employed. The previous study typically identified a mixed population of senescent cells, macrophages, FAPs and MuSCs, using SA-β-gal staining, and subsequently utilized p16^INK4a^-based transgenic models (*p16-3MR* mice) or broad-spectrum senolytics like Dasatinib and Quercetin (D+Q) or ABT-263 to effect global clearance. However, our current study and others demonstrate *Cdkn2a* (p16^INK4a^) is expressed at exceedingly low levels in senescent MuSCs ^18^, making it an inadequate marker for this lineage. It is therefore highly probable that the regenerative impairments observed in Moiseeva’s study were a consequence of clearing senescent macrophages or FAPs, which may indeed harbor detrimental functions. Furthermore, the *p16-3MR* model has been shown to exhibit inefficient clearance of senescent cells, thereby confounding data interpretation^24^. In contrast, our study focused specifically on the newly identified, senescence-like USS^Hi^ MuSC subpopulation. By utilizing a high-resolution *Pax7-CreERT; Ccl2-LSL-DTA* genetic model driven by the specific marker *Ccl2*, we were able to selectively ablate this specialized population while leaving other niche constituents and the normal USS^Lo^ MuSC pool unperturbed. This lineage-specific approach provides solid genetic evidence to show that senescence-like USS^Hi^ MuSCs are essential, pro-regenerative components of the early injury response.

Adding to the complex nature of the cells, MHCII surface marker was also used for defining “immunomyoblasts”, a population of MuSCs primarily characterized during the mid-to-late stages of skeletal muscle regeneration. Like USS^Hi^ MuSCs, IMBs were also characterized by the robust expression of immune-related genes^26^. Again, lineage tracing will be necessary to examine whether the two populations are from the same origin or the early emergence of the USS^Hi^ cells may prime the injury niche prior to the onset of IMBs.

### Multiomics mapping and spatial transcriptomics validate the USS^Hi^ MuSC-dependent niche network in regenerating muscle

The mechanistic basis for this pro-regenerative function lies in the extraordinary secretory capacity of the USS^Hi^ MuSCs. The initial phase of muscle repair is featured by complex and hierarchical events of fiber necrosis, immune cell influx and the subsequent transition from an inflammatory to a reparative, myogenic phase. ^43,44^. In the current study, our matched transcriptomic, proteomic, and secretomic profiles comprehensively demonstrate that USS^Hi^ cells represent a unique subpopulation of MuSCs that undergo intracellular reprogramming to robustly express and secrete classical pro-inflammatory factors, including CCL2, CCL3, CXCL2, and IL1b etc. Further integration of high-resolution spatial transcriptomics and scRNA-seq elucidates the specific cellular recipients of these signals and the consequent spatiotemporal alterations in regenerating tissue architecture. By projecting segmented cellular coordinates onto the transcriptomic landscape of the injury zone, we demonstrated that the loss of USS^Hi^ MuSCs both physically distances the stem cells from their neighboring niche cells and cuts off the signaling between them. The ablation of this minor population led to a collapse in paracrine communication networks, fundamentally disrupting critical interactions with myeloid cells, FAPs, and ECs. These findings reinforce the concept that USS^Hi^ MuSCs arising in the early injury milieu function as crucial signaling hubs, actively orchestrating immune cell recruitment, polarization and stromal activation as well as EC expansion/angiogenesis within the injury niche via their localized secretory phenotype.

### Bioactive lipid S1P mediated dual-axis mechanism driving angiogenesis and MuSC expansion in regenerating muscle

The transcriptomic interaction inference relies exclusively on protein-coding ligand-receptor pairs, rendering it inherently blind to lipid-and metabolite-mediated signaling. Indeed, the functional roles of lipid SASP components remain largely understudied. While the accumulation of intracellular LDs is a frequently observed morphological feature of senescent cells across various tissues^45^, the functional consequences of this lipid dysregulation have rarely been extended to intercellular communication. Importantly, these LDs are not metabolically isolated entities. It is likely that as the lipid-buffering capacity of LDs reaches saturation in USS^Hi^ MuSCs, excess lipid flux is shunted toward sphingolipid metabolism^46^. Therefore, we conducted the characterization of lipid-mediated secretome driven by metabolic rewiring within the USS^Hi^ MuSCs. Indeed, our results revealed that USS^Hi^ MuSCs do not merely store neutral lipids as a passive reservoir; rather, this metabolic rewiring drives a highly coordinated, unidirectional catabolism of complex glycosphingolipids, such as LacCer and GluCer. The concerted upregulation of distinct lysosomal and cytosolic enzymes (GLB1, GBA1/2, ASAH1/2, SPHK1/2) creates a metabolic funnel that couples this intracellular lipid overflow to the terminal synthesis and subsequent extracellular release of the bioactive lipid S1P. It is conceivable that S1P is packaged within extracellular vesicles (EVs) to protect it from rapid degradation and facilitate its targeted delivery to the surrounding niche, representing an intriguing avenue for future investigation.

Although S1P is widely recognized for its systemic functions, including the regulation of lymphocyte trafficking, the preservation of vascular integrity, and the modulation of cytokine and chemokine networks that sustain immune and cardiovascular homeostasis^47^, our study reveals an important role for this lipid mediator in orchestrating local muscle repair.

Specifically, we demonstrate that the localized secretion of S1P by USS^Hi^ MuSCs establishes a highly specific, dual-axis signaling network within the regenerative microenvironment. Our single-cell network analysis, supported by *in vitro* functional assays, *in vivo* genetic evidence and pharmacological inhibition, delineates the divergent downstream targets of this lipid-driven signaling axis. Firstly, S1P engages the S1PR1 receptor exclusively expressed on the local microvasculature, driving the proliferation and migration of ECs to ensure adequate angiogenesis. Concurrently, S1P acts in a paracrine manner upon adjacent, typical USS^Lo^ MuSCs via the S1PR2 receptor to stimulate their robust proliferation, thereby driving the massive cellular expansion necessary for timely myofiber repair. The profound depletion of CD31+ ECs following USS^Hi^ MuSC ablation *in vivo* highlights the dependency of the vascular niche on USS^Hi^ MuSCs-derived S1P. These findings are substantiated by our *in vivo* rescue experiments, where exogenous S1P administration was sufficient to bypass the ablation of USS^Hi^ MuSCs, successfully restoring the PAX7^+^ stem cell pool, re-establishing the vascular network, and rescuing myofiber maturation. Although S1P can be ubiquitously produced by various local cell types, including tenocytes and adjacent endothelial cells, our ablation data demonstrate that these alternative sources fail to functionally compensate for the depletion of USS^Hi^ MuSCs (Fig. 4 and 5). These data position S1P as the critical molecular bridge linking USS^Hi^ MuSC metabolic remodeling to holistic tissue regeneration.

In summary, our study redefines the functional boundaries of cellular senescence within the context of skeletal muscle regeneration. By isolating the distinct USS^Hi^ subpopulation and uncovering a novel sphingolipid-mediated communication axis, we demonstrate that a senescence-like state in stem cells is a meticulously programmed, requisite driver of early tissue repair. Additional to a classical protein-dominated secretory profile the recognition of a functionally decisive lipid-mediated secretome reshapes our mechanistic understanding of the regenerative niche and highlights targeted lipidomic signaling as a highly promising frontier for future regenerative medicine. While current findings demonstrate that these cells are indispensable for initiating the repair cascade during this early period, the precise mechanisms underlying their emergence remain to be fully elucidated. Given their validated inflammatory phenotype and rapid emergence following acute injury, we speculate that stimuli from the extrinsic inflammatory niche induce a subset of activating MuSCs to exhibit inflammatory characteristics, thereby generating the USS^Hi^ population. Further study is needed to test this hypothesis. Additionally, the temporal dynamics and functional roles of these cells in regenerating muscle of aged mice could also be tested; the emergence timeline and functional consequence may be dramatically different from what we observed in young mice due to fundamentally altered niche milieu. It is highly plausible that while the USS^Hi^ state is a beneficial, acute driver of repair in young tissue, the age-related persistence or aberrant terminal fate of these cells could contribute to a chronic pro-inflammatory and pro-fibrotic environment that inherently restricts regenerative capacity.

## Resource Availability Lead contact

Requests for further information and resources should be directed to and will be fulfilled by the lead contact, Huating Wang.

## Materials availability

All reagents generated in this study are available from the lead contact upon completion of a material transfer agreement.

## Data and code availability

Protein mass spectrometry data have been deposited to ProteomeXchange (http://www.proteomexchange.org) with the dataset identifier PXD082531 and PXD082720. The lipidomics data generated in this study are provided in the Source Data file. Bulk RNA-seq, single-cell RNA-seq and spatial transcriptomics data have been submitted to Gene Expression Omnibus (https://www.ncbi.nlm.nih.gov/geo/) with the accession code GSE344130, GSE344176 and GSE344178. This paper does not report any original code. All other data supporting the findings of this study are available from the corresponding author on reasonable request.

## Supporting information

Introduction of supplementary files

## Acknowledgments

This work was supported by Non-Communicable Chronic Disease-National Science and Technology Major Project of China to H.W. (project code: 2024ZD0530400); National Key R&D Program of China to H.W. (project code: 2022YFA0806003); The research funds from Health@InnoHK program launched by Innovation Technology Commission, the Government of HK to H.W.; Health and Medical Research Fund (HMRF) from Health Bureau of HK to H.W. (project codes: 10210906 and 08190626); Theme-based Research Scheme (TRS) from RGC (project code:T13-602/21-N); General Research Fund (GRF) from Research Grants Council (RGC) of the Hong Kong (HK) Special Administrative Region, China to H.W. (project codes: 14105123, 14103522, 14105823, 14108225 and 14103526 to H.W.); the National Natural Science Foundation of China (NSFC) to H.W. (project codes: 82172436 and 31871304); Area of Excellence Scheme (AoE) from RGC to H.W. (project code: AoE/M-402/20); 1+1+1 CUHK-CUHK(SZ)-GDST Joint Collaboration Fund to H.W (project code: 524002645); Strategic Topics Grant (STG) from RGC to H.W. (project code: STG1/M-404/26-N).

## Author contributions

X.L. performed the wet-lab experiments, multiomics data analysis and single-cell RNA-seq analysis; Y.Z. analyzed the spatial transcriptomics data; Y.L., Y.J. and Y.Q. performed and helped with wet-lab experiments and single-cell RNA-seq; C.L. helped with the single-cell RNA-seq analysis; S.W. and C.C.L.W. helped and performed the secretome; S.M.L. and G.S. helped and performed the proteome; H.S. supervised computational analyses; S.K. guided this project and helped with the USS^Hi^ MuSCs sorting strategy; H.W. supervised experiments; X.L. and H.W. conceived the project and wrote the manuscript, with inputs from all authors.

## Declaration of interests

No potential conflict of interest relevant to this article was reported.

## Declaration of generative AI and AI-assisted technologies in the manuscript preparation process

During the preparation of this work, the authors used Gemini and Kimi to assist with grammar and language refinement. After using these tools and suggestions, the authors reviewed and edited all content as needed and took full responsibility for the content of the published article.

## Methods

### Mice

*Pax7-CreER (Pax7tm1(cre/ERT2)Gaka)*, *Pax7-nGFP* and *ROSAEYFP* mouse strains were kindly provided by Dr. Zhenguo WU (Hong Kong University of Science and Technology). *Ccl2-LSL-DTA* knock-in mice were generated by GemPharmatech (Nanjing, China). *Pax7-CreER* mice were crossed with *ROSAEYFP* mice to generate the *Pax7-CreER; ROSAEYFP* reporter mice. The Ctrl (*Pax7-CreER+/+; ROSAEYFP+/+; Ccl2-LSL-DTA-/-*) and KI (*Pax7-CreER+/+; ROSAEYFP+/+; Ccl2-LSL-DTA+/+*) mice were generated by crossing *Ccl2-LSL-DTA* mice with *Pax7-CreER; ROSAEYFP* reporter mice. Primers used for genotyping are shown in Suppl. Dataset 10. The mice were maintained in the animal room with 12h light/12h dark cycles, 22–24 °C room temperature, and 40–60% humidity at the animal facility at the Chinese University of Hong Kong (CUHK). All animal handling procedures and protocols were approved by the Animal Experimentation Ethics Committee (AEEC) of CUHK (Ref. No. 23-195-MIS). All animal experiments followed the regulations and guidance of laboratory animals of CUHK.

### Animal procedures

Inducible specific ablation of USS^Hi^ MuSCs was administered by injecting Tamoxifen (Tmx) (T5648, Sigma) intraperitoneally (IP) at 2 mg per 20 g body weight. For BaCl_2_-induced muscle injury, 2-3-month-old mice were intramuscularly injected with 50 μl of 1.2% BaCl_2_ (B861682, MACKLIN) solution into TA muscles, and the muscles were harvested at designated time points for further analysis. For S1P treatment in vivo, 50 μl 500 μM S1P (HY-108496, MCE) was injected intramuscularly into TA muscles on the second day after BaCl_2_ injection for 3 days, and the muscles were harvested at designated time points for further analysis.

### Fluorescence-activated cell sorting

Following our established protocol^48^, injured or uninjured hindlimb muscles from *Pax7-nGFP*, Ctrl and KI mice were dissected and minced with blades, then digested with collagenase II (1100 U ml-1, Worthington) in Ham’s F-10 media (Thermo Fisher Scientific) for 90 min at 37°C with gentle rotation at 70 rpm. The digested muscles were washed in the washing medium (Ham’s F-10 media, 10% HIHS (Gibco), penicillin/streptomycin (1×, Gibco)) once, and MuSCs were further released by treating muscles with Collagenase II (1100 U ml-1) and Dispase (11 U ml-1) for 40 min at 37°C. Digested tissue was passed through a 21-gauge needle 12 times and filtered through a 40-μm filter followed by spinning at 700 × g for 5 min at 4 °C. Mononuclear cells were resuspended and filtered with a 40-µm cell strainer, and GFP+/EYFP+ MuSCs or CD45-CD31+ muscle endothelial cells were sorted out by BD FACS Aria Fusion cell sorter (BD Biosciences).

### Cell culture

Freshly isolated MuSCs were cultured in Ham’s F10 medium supplemented with 20% FBS (growth medium) on dish/slides precoated with PDL and ECM. Freshly isolated muscle endothelial cells were cultured in EGM®-2 Endothelial Cell Growth Medium-2 BulletKit (CC-3162, Lonza). Mouse MS1 endothelial cells (CRL-2279) were kindly provided by Dr. Kathy LUI and cultured in DMEM medium with 5% fetal bovine serum, 100 units/ml of penicillin, and 100 μg of streptomycin (growth medium, or GM) at 37 °C in 5% CO2 incubator. For S1P treatment, cells were exposed to S1P (MCE, HY 108496) at a final concentration of 2 μM for 24 h. For JTE 013 treatment, cells were treated with JTE 013 (MCE, HY 100675) at a final concentration of 10 μM for 48 h. For W146 treatment, cells were incubated with W146 (MCE, HY 101395) at a final concentration of 10 μM for 27 h.

### EdU assay

EdU assay was performed following our protocol^49^ using the Invitrogen™, Click-iT™ EdU Cell Proliferation Kit for Imaging, Alexa Fluor™ 594 dye (C10339). MS1 cell line was incubated with 10 μM EdU for 4hr before fixation with 4% PFA. Isolated MuSCs and primary endothelial cells were incubated with 10 μM EdU for 24 hr before fixation with 4% PFA.

### S1P ELISA

FACS-isolated USS^Lo^ and USS^Hi^ MuSCs (collected at 3 dpi) were plated at equal densities and cultured in serum-free medium for 24 h. The conditioned medium was then collected and centrifuged at 2,000 × g for 10 min at 4 °C to remove cells and debris. S1P concentration in the supernatant was measured using a commercial competitive S1P ELISA kit (FineTest, Cat. No. EU2603) according to the manufacturer’s instructions. Briefly, samples and serially diluted S1P standards were added to pre-coated wells, followed by sequential incubation with detection reagents and substrate solution. Absorbance was measured at 450 nm using a microplate reader (CLARIOstar Plus, BMG LABTECH). S1P concentrations were interpolated from the standard curve using four-parameter logistic regression and normalized to the input cell number.

### Immunofluorescence, immunohistochemistry and lipid droplets staining

For immunofluorescence staining, cultured cells were fixed in 4% PFA for 15 min and permeabilized with 0.5% NP-40 for 10 min. Then cells were blocked in 3% BSA for 1 h followed by incubating with primary antibodies (PAX7 (Developmental Studies Hybridoma Bank; 1:30) and S1PR2 (Thermo Scientific, PA523208; 1:200)) overnight at 4 °C and secondary antibodies for 1 hr at RT. Finally, the cells were mounted with DAPI to stain the cell nucleus and images were captured by a Leica fluorescence microscope. For immunohistochemistry, in brief, slides were fixed with 4% PFA for 15 min at room temperature and permeabilized in ice-cold methanol for 6 min at −20 °C. Heat-mediated antigen retrieval with a 0.01 M citric acid (pH 6.0) was performed for 5 min in a microwave. After 4% BSA (4% IgG-free BSA in PBS; Jackson, 001-000-162) blocking, the sections were further blocked with unconjugated AffiniPure Fab Fragment (1:100 in PBS; Jackson, 115-007-003) for 30 min. The biotin-conjugated anti-mouse IgG (1:500 in 4% BSA in PBS, Jackson, 115-065-205) and Cy3-Streptavidin (1:1250 in 4% BBBSA, Jackson, 016-160-084) were used as secondary antibodies. Primary antibodies and dilutions were used as follows: CD31 (Thermo, 14-0311-82; 1:500), PAX7 (Developmental Studies Hybridoma Bank; 1:30), MYOD (Dako, M3512; 1:500), eMyHC (Leica NCL-MHC-d; 1:200), Laminin (Sigma-Aldrich L9393-100UL, 1:800) for staining of muscle cryosections. Images were slightly modified with ImageJ in which background was reduced using background subtraction and brightness and contrast were adjusted. H&E (Hematoxylin-eosin) staining on TA muscle sections was performed according to protocol. Section slides were first stained in hematoxylin for 10 min followed by rinsing thoroughly under running tap water for at least 3 min. Then section slides were immersed in 0.2% acid alcohol for 1 s and immediately rinsed under running tap water. Next, section slides were stained in eosin for 2 min followed by rinsing and dehydrating in graded ethanol and Xylene. Finally, slides were mounted by DPX and observed under a normal microscope. Lipid droplets were visualized using the Lipid Droplets Green Fluorescence Assay Kit (Beyotime, C2053S) according to the manufacturer’s instructions.

### SA-β-galactosidase staining

Cellular senescence was evaluated by β-galactosidase activity using β-galactosidase Senescence Kit (#9860, Cell Signaling Technology)^12^. Briefly, cells were fixed for 15 min followed by washing in PBS twice. Then fixed cells were incubated with β-galactosidase staining solution at 37 °C in a dry incubator (no CO2) at least overnight. The cells were then observed under a microscope for the development of blue color.

### RNA isolation and quantitative RT-PCR

Total RNAs were extracted using TRIzol reagent (Invitrogen) following the manufacturer’s protocol. For quantitative RT-PCR, cDNAs were reverse transcribed using HiScript III 1st Strand cDNA Synthesis Kit (Vazyme, R312-01). Sequences of all primers used can be found in Suppl. Dataset 10. Real-time PCR reactions were performed on a LightCycler® 480 Instrument II (Roche Life Science) using Luna Universal qPCR Master Mix (NEB, M3003L).

### Bulk RNA-seq and data analysis

For bulk RNA-seq (polyA+ mRNA), total RNAs were subject to polyA selection (Ambion, 61006) followed by library preparation using NEBNext® Ultra™ II RNA Library Preparation Kit (NEB, E7770S). Libraries were paired-end sequenced with read lengths of 150 bp on Illumina HiSeq X Ten or Nova-seq instruments. The raw reads of RNA-seq were processed following the procedures described in our previous publication. Briefly, the adapter and low-quality sequences were trimmed from 3’ to 5’ ends for each read, and the reads shorter than 50 bp were discarded. The clean reads were aligned to the mouse (mm10) reference genome with STAR. Next, we used DESeq2 to quantify the gene expression. Genes were identified as differentially expressed genes (DEGs) if the change of expression level is greater than two folds and the p-value is less than 0.05 between two stages/conditions. GO enrichment analysis was performed using the R package *clusterProfiler*.

### Single-cell RNA-seq and data analysis

Single-cell RNA-seq was performed on 10x genomics platform following our established protocol^44^. Briefly, MuSCs or whole muscle cells were isolated with an additional step of viability validation by Propidium Iodide (PI) staining. Red blood cells were eliminated by ACK buffers (150 mM NH4Cl, 100 mM KHCO3, 10 mM EDTA-2Na) before sorting. After sorting, live cells were washed with 0.04% BSA in PBS twice and resuspended in the BSA solution at an appropriate concentration (300– 1200 cells/μl). Suspended cells were counted under a microscope and Trypan blue was used to examine cell viability. After the confirmation of cell number and viability, library construction was performed following the manufacturer’s instructions for generation of Gel Bead-In Emulsions (GEMs) using the 10x Chromium system. The single cell RNA library was sequenced by Illumina HiSeq X Ten instrument. CellRanger (v8.0.0) and Seurat (v5.4.0)^50^ were used to analyze the single-cell data. Samples were initially merged together and filtered with quality control parameters (cells with more than 10% expression on mitochondrial genes, or fewer than 500 total features expressed were filtered out). Replicates and samples were integrated using Harmony within the Seurat framework to mitigate batch effects.

### Cell fate trajectory analysis

Pseudotime trajectory analysis was performed using Monocle 2 (v2.14.0)^51^ on a downsampled dataset of 10,000 cells. RNA count data from the Seurat sketch assay was extracted and used to construct a CellDataSet object. Ordering genes were selected from USS gene set^12^ based on mean expression ≥ 0.1 using *setOrderingFilter*. Dimensionality reduction was performed using the DDRTree method. Pseudotime ordering was calculated, with the root state defined as the state containing the maximum number of 0 dpi cells. The cells falling into late 1 and late 2 branches were extracted, and DEGs between the two late fates were detected by *FindMarkers*. To quantify cell type composition changes along the pseudotime trajectory, pseudotime values were binned into discrete intervals of 1 unit using the cut function. For each cell type, the proportion of cells within each pseudotime bin was calculated. The relationship between pseudotime bin and cell type proportion was assessed using Spearman correlation (cor.test, method = “spearman”). Scatter plots with linear regression lines were generated using ggplot2, with correlation coefficients and P-values annotated on each plot.

### Senescent cell identification by Unified Senescence Score

Unified Senescence Score method was performed as previously described with minor modifications^12^. Briefly, four established senescence-associated gene databases were integrated: SenMayo (SM), CellAge (CA), GenAge (GA), and Senescence Eigengene (SE). Known senescent cell markers (p16^INK4a^, p15, p19, p21, p27, and PAI-1) were excluded from the combined gene set. The USS score was calculated using the *AddModuleScore* function in Seurat. Cells were classified as USS^Hi^ when their USS score exceeded 0.1745.

### Differential expression and Gene Ontology enrichment analysis

Differentially expressed genes (DEGs) were analyzed by *FindMarkers* function in Seurat using Wilcoxon Rank Sum test and were detected with the cutoff of Log2FC >0.5 and adjusted *P*-value <0.05. Gene Ontology (GO) enrichment analysis for DEG was performed with the *enrichGO* function in the *clusterProfiler* (version 4.2.0) package. GO biological terms with Benjamini-Hochberg adjusted *P*-value (FDR) < 0.05 were considered significantly enriched.

### Cell-cell communication analysis

Cell-cell interactions were inferred by CellChat (version 2.2.0)^52^ based on the expression of known ligand-receptor pairs in various cell types. Cells from Ctrl and KI groups were applied to CellChat separately and merged into one CellChat object for comparison analysis following CellChat tutorial.

### Visium HD spatial transcriptomics

For tissue section preparation, the target muscles were extracted from the mice and placed in 30% sucrose at 4 for dehydration. They were then embedded with OCT and transported on dry ice, and quickly frozen and stored at-80. At this point, the sample freezing process was completed. When slicing was needed, the dry ice was taken out from-80 and transported to the slicing machine for slicing. Each tissue slice had a thickness of 10 micrometers. 20 slices were collected for RNA quality detection. The RNA integrity number (RIN) value must be greater than 7 for the sample to be used; one slice was placed on a glass slide for staining, and only samples with normal pathological structures could undergo the formal experiment. After passing the quality inspection, the samples were selected for sectioning. The slices expressing the target location (each 10 µm thick) are used for the formal experiment. These slices are prepared, fixed and stained according to the Visium HD Fresh Frozen Tissue Preparation Handbook (CG000763). Fluorescent microscope (Pannoramic MIDI, 3DHISTECH) is used for imaging. Destain and permeabilization was then performed, followed by probe hybridization, probe ligation, Visium HD slide preparation, probe release, extension, and library construction using the Visium HD Spatial Gene Expression Reagent Kits User Guide (CG000685). The final libraries were quantified using the Qubit High Sensitivity DNA assay (Thermo Fisher Scientific) and the size distribution of the libraries were determined using a High Sensitivity DNA chip on a Bioanalyzer 4150 (Agilent). Sequencing was performed on an DNBSEQ-T7 Sequencer (MGI, Shenzhen, China) with paired-end reads.

### Spatial transcriptomics data analysis

After sequencing, raw FASTQ files were aligned to the mouse reference transcriptome (mm10-2020-A) using the 10x Genomics Space Ranger (v4.0.1) count function. Space Ranger generated Visium HD data binned at 2-μm, 8-μm, and 16-μm resolutions, along with nuclear segmentation results. Downstream analyses were performed on both the nuclear segmentation and 8-μm resolution data. Regions with distinct injury levels (injury, peri-injury and non-injury) were manually annotated on the 8-μm resolution data based on H&E staining images using Loupe Browser. These injury level annotations were subsequently transferred to the nuclear segmentation data based on spatial coordinates.

For the nuclear segmentation data, analysis was conducted using Seurat (v5.3.1, R v4.4). Cells were filtered based on mitochondrial content (< 5%) and UMI/gene counts (2.5-97.5 percentile range) to exclude low-quality cells and outliers. Data were normalized using *NormalizeData*, and highly variable features were identified using *FindVariableFeatures*. Following scaling with *ScaleData*, Principal Component Analysis was performed using *RunPCA*. Samples were then integrated using *IntegrateLayers* with the RPCA integration method (method = RPCAIntegration, orig.reduction = “pca”). After integration, layers were re-joined using *JoinLayers*. Cell clustering was performed using *FindNeighbors* with the integrated PCA coordinates (reduction = “integrated.cca”, dims = 1:30), followed by *FindClusters*. Finally, UMAP dimensionality reduction was performed using *RunUMAP* with 30 principal components from the integrated reduction for visualization. Cell types were annotated based on canonical marker gene expression.

For cell-cell communication analysis, cells from injury level regions were extracted for cell-cell interaction analysis using CellChat (v2.1.2). For each sample, the two slides were designated as biological replicates. Cell-cell communication probabilities were computed using *computeCommunProb* with the following parameters: type = “truncatedMean”, trim = 0.1, distance.use = TRUE, interaction.range = 500, scale.distance = 0.5, contact.dependent = TRUE, and contact.range = 10. Muscle injury regeneration-specific signaling pathways and ligand-receptor pairs were identified and visualized using built-in functions in CellChat.

Spatial cell type co-occurrence analysis was performed on cells from injury regions using Squidpy^53^. A spatial neighborhood graph was constructed using *sq.gr.spatial_neighbors* with a 100 μm radius (coord_type = “generic”). Co-occurrence probabilities were calculated using sq.gr.co_occurrence across increasing radii from 1 to 200 μm (interval = 1 μm steps).

### Secretomics and data analysis

The 1 dpi and 3 dpi MuSCs were isolated by FACS and cultured with serum-free Ham’s F-10 Nutrient Mixture for 24 hours to collect secreted proteins. The supernatant was then centrifuged at 2000g for 10 min to remove cell debris. Following protein quantification of the culture medium samples via the bicinchoninic acid assay, an appropriate volume of each sample was mixed with ice-cold acetone and incubated at −20 ° C for several hours to precipitate proteins. The precipitate was collected by centrifugation at 18,000 × g for 30 min at 4°C, and the supernatant was discarded. The resulting protein pellet was resuspended and thoroughly mixed in a denaturation buffer containing 8 M urea and 100 mM Tris-HCl (pH 8.5). For reduction and alkylation, tris-(2-carboxyethyl)-phosphine (TCEP) was added to a final concentration of 5 mM and incubated at room temperature for 30 min, followed by the addition of iodoacetamide (IAA) to a final concentration of 10 mM and incubation in the dark for an additional 30 min. For enzymatic digestion, the sample was diluted with a dilution buffer (100 mM Tris-HCl, pH 8.5) and thoroughly vortexed. Trypsin (1 μg) was subsequently added, and the mixture was incubated in a dry block heater at 37°C for 18 h. Finally, the digested peptides were desalted using a desalting column.

For liquid chromatography, a nanoElute liquid chromatography system (Bruker Daltonics) was applied for peptide separation at a flow rate of 300 nL/min. Water and ACN with 0.1% formic acid were mobile phase A and B, respectively. A nonlinear gradient separation process was set to firstly increase from B% at 2%–22% within the time range of 40 min, secondly increase from B% at 22%–37% within 10 min, thirdly increase from B% at 37% – 100% within 5 min, and finally maintained at 100% for the last 5 min before re-equilibration.

For mass spectrometry, all samples above were analyzed on timsTOF Pro2 (Bruker). A CaptiveSpray nanoelectrospray ion source was applied to interface LC with mass spectrometer. Generation of the spectral library was operated in data-dependent mode, with the accumulation and ramp time ranging from m/z 100 to 1700 in positive electrospray mode. The sample analysis was performed in data-independent mode, with an MS1 scan and 64 MS2 windows in a diaPASEF acquisition scheme covering the mass range from m/z 400 to m/z 1200. During the scanning mode, the ion mobility was set from 0.6 to 1.6 Vs/cm2, and the collision energy was descended linearly as a function of the mobility from 59 eV at 1/K0 = 1.6 Vs/cm2 to 20 eV at 1/K0 = 0.6 Vs/cm. Spectral libraries were generated with Spectronaut version 19.2 (Biognosys) against a UniProt Mouse database (accessed on June 27, 2024, only reviewed entries). All the parameters were default.

### Proteomics and data analysis

3 dpi MuSCs were isolated by FACS and suspended in Methanol solution. For protein extraction, each sample mixed with 8 µL of 50 mM ammonium bicarbonate (Sigma) and 100 ng trypsin, mixed by pipetting up and down, and centrifuged at 1000g for 2 min, followed by incubation at 37 °C for 6 h. Digestion was stopped by adding formic acid (FA) to a final concentration of 1% (v/v). The samples were then centrifuged at 13,000 × g for 10 min at room temperature, and the supernatant was collected for LC–MS/MS analysis.

The peptides were reconstituted in buffer A (0.1% formic acid water) and separated on a 25 cm analytical column with an integrated emitter tip (75 µm ID, 1.6 µm C18 beads, Aurora Series with CSI, IonOpticks, Australia) maintained at 50 °C in a thermostated column oven. Samples were analyzed on nanoliter UHPLC (Thermofisher, USA) and timsTOF Pro2 (Bruker Daltonics, Germany) equipped with a nanoliter spray ion source. Samples were analyzed at a flow rate of 300 nL/min: buffer B increased from 6 to 30% at 22 min, to 40% at 6 min, to 99% at 0.1 min, and remained 99% at 3 min. For the MS setup, timsTOF Pro2 operates in data-independent acquisition combined with parallel accumulation-serial fragmentation (dia-PASEF) mode. The capillary voltage was set to 1600 V, the drying temperature was 180, the flow rate was 3.0 L/min, the primary and secondary mass spectrum collection range was 100-1700 m/z, and the ion mobility range (1/K0) was 0.6-1.6 Vs/cm2. The collision energy range is 20-59 eV with 32 windows for acquisition^54^.

For database search, raw data was searched against a preferred database using Spectronaut 19.0 (Biognosys). The digestion enzyme was Trypsin/P and the max missed cleavage was set at 2. The methionine oxidation and acetylation of the protein N-terminus were set as variable modifications. FDR was controlled to 1% at both protein and peptide level. The minimum one peptide per protein was allowed for protein identification. The rest was set as default^55^. The mouse database (17212 sequences) was downloaded from UniProt on August 06, 2024.

### Lipidomics and data analysis

3 dpi MuSCs were isolated by FACS and suspended in Methanol solution. For lipid extraction, lipids were extracted from cells vesicles using a modified version of the Bligh and Dyer’s method as described previously^56^. Briefly, cells were homogenized in 750 µL of chloroform: methanol: MilliQ H2O (3:6:1) (v/v/v). The homogenate was then incubated at 1500 rpm for 1h at 4. At the end of the incubation, 350 µL of deionized water and 250 µL of chloroform were added to induce phase separation. The samples were then centrifuged and the lower organic phase containing lipids was extracted into a clean tube. Lipid extraction was repeated once by adding 450 µL of chloroform to the remaining cells in aqueous phase, and the lipid extracts were pooled into a single tube and dried in the SpeedVac under OH mode. Samples were stored at-80 until further analysis.

Lipidomic analyses were conducted at LipidALL Technologies using a ExionLC-AE coupled with Sciex QTRAP 7500 PLUS as reported previously^57^. Separation of individual lipid classes of polar lipids by normal phase (NP)-HPLC was carried out using a TUP-HB silica column (i.d. 150×2.1 mm, 3 µm) with the following conditions: mobile phase A (chloroform: methanol: ammonium hydroxide, 89.5:10:0.5) and mobile phase B (chloroform: methanol: ammonium hydroxide: water, 55:39:0.5:5.5). MRM transitions were set up for comparative analysis of various polar lipids. Individual lipid species were quantified by referencing to spiked internal standards. d9-PC32:0(16:0/16:0), d9-PC36:1p(18:0p/18:1), d7-PE33:1(15:0/18:1), d9-PE36:1p(18:0p/18:1), d31-PS(d31-16:0/18:1), d7-PA33:1(15:0/18:1), d7-PG33:1(15:0/18:1), d7-PI33:1(15:0/18:1), C17-SL, d5-CL72:8(18:2)4, Cer d18:1/15:0-d7, d9-SM d18:1/18:1, C8-GluCer, C8-GalCer, d3-LacCer d18:1/16:0, Gb3 d18:1/17:0, d7-LPC18:1, d7-LPE18:1, C17-LPI, C17-LPA, C17-LPS, C17-LPG, d17:1-Sph were obtained from Avanti Polar Lipids. GM3-d18:1/18:0-d3 was purchased from Matreya LLC. Free fatty acids were quantitated using d31-16:0 (Sigma-Aldrich) and d8-20:4 (Cayman Chemicals).

Glycerol lipids including diacylglycerols (DAG) and triacylglycerols (TAG) were quantified using a modified version of reverse phase HPLC/MRM^58^. Separation of neutral lipids was achieved on a Phenomenex Kinetex-C18 column (i.d. 4.6×100 mm, 2.6 µm) using an isocratic mobile phase containing chloroform:methanol:0.1 M ammonium acetate 100:100:4 (v/v/v) at a flow rate of 300 µL for 10 min. Levels of short-, medium-, and long-chain TAGs were calculated by referencing to spiked internal standards of TAG (14:0)3-d5, TAG (16:0)3-d5 and TAG (18:0)3-d5 obtained from CDN isotopes, respectively. DAGs were quantified using d5-DAG 17:0/17:0 and d5-DAG18:1/18:1 as internal standards (Avanti Polar Lipids).

Free cholesterols and cholesteryl esters were analyzed under atmospheric pressure chemical ionization (APCI) mode on a Jasper HPLC coupled to Sciex 4500 MD as described previously, using d6-cholesterol and d6-C18:0 cholesteryl ester (CE) (CDN isotopes) as internal standards^59^.

### Statistical and reproducibility

Statistical tests were performed for independent-samples with unpaired *t*-test, paired *t*-test or one-way ANOVA tests (GraphPad Prism, version 8; GraphPad Software, San Diego, CA). All statistical tests incorporated two-tailed tests and homogeneity of variance tests and were considered to reflect significant differences if \**P* < 0.05, \*\**P* < 0.01, \*\*\**P* < 0.001 or \*\*\*\**P* < 0.0001. Data are represented as the mean of at least three biologically independent samples ± SD. Details of statistical analyses including sample numbers (*n*) are included in the respective figure legends.

## Notes

### Competing Interest Statement

The authors have declared no competing interest.

