## Supplementary material for "Multiomics Characterization Identifies S1P-secreting USS^High^ Skeletal Muscle Stem Cells as Essential Drivers of Niche Remodeling and Muscle Regeneration": Introduction of supplementary files

**Supplementary Materials**

**List of Supplementary Figures**

Supplementary Figure S1. Characterization of MuSC subclusters and pseudotime dynamics during acute injury induced early regeneration.

Supplementary Figure S2. FACS gating strategy for MHC-II-based isolation of USS^Hi^ MuSCs.

Supplementary Figure S3. Secretomic and lipidomic analyses of USS^Hi^ and USS^Lo^ MuSCs.

Supplementary Figure S4. Single-cell validation of the Ccl2-DTA ablation model.

Supplementary Figure S5. Extended spatial and single-cell transcriptomics analyses of the regenerating niche.

Supplementary Figure S6. FACS gating strategy for EC isolation from wild type mice skeletal muscle at 3 dpi.

Supplementary Figure S7. Baseline MuSC abundance in Ctrl and KI mice.

**List of Supplementary Tables**

Supplementary Data 1. scRNA-seq analysis of isolated MuSCs.

Supplementary Data 2. Bulk RNA-seq analysis of isolated USS^Hi^ and ^Lo^ MuSCs.

Supplementary Data 3. Proteomic analysis of USS^Hi^ and USS^Lo^ MuSCs.

Supplementary Data 4. Secretomic analysis of USS^Hi^ and USS^Lo^ MuSCs.

Supplementary Data 5. Integrated analysis of proteomics and secretomics of USS^Hi^ and USS^Lo^ MuSCs.

Supplementary Data 6. Lipidomic analysis of USS^Hi^ and USS^Lo^ MuSCs.

Supplementary Data 7. scRNA-seq profiling of MuSCs in Ctrl and KI muscles at 3 dpi.

Supplementary Data 8. Spatial transcriptomic analysis of Ctrl and KI muscles at 3 dpi.

Supplementary Data 9. scRNA-seq profiling of Ctrl and KI muscles at 1 and 3 dpi.

Supplementary Data 10. Sequences of oligonucleotides used in the study.

**Supplementary Figure legend**

**Supplementary Figure S1. Characterization of MuSC subclusters and pseudotime dynamics during acute injury induced early regeneration. (A)** Violin plots showing the expression of canonical cell identity markers across the nine MuSC subclusters at 0, 1 and 3 dpi. **(B)** Percentage of each MuSC subpopulation along the pseudotime of the Late 1 and Late 2 branches, with Spearman correlation coefficients (R) and P values. **(C)** Dot plot showing the percentage of USS^Hi^ cells and the total cell number of each subpopulation across 0, 1 and 3 dpi.

**Supplementary Figure S2. FACS gating strategy for MHC-II-based isolation of USS^Hi^ MuSCs. (A)** Representative plots showing sequential gating strategy: FSC-A/SSC-A for bulk cells, FSC-H/FSC-A and SSC-H/SSC-A for doublet exclusion, and PE-A (MHCII) for the separation of MHCII+ (USS^Hi^) from MHCII- (USS^Lo^) MuSCs.

**Supplementary Figure S3. Secretomic and lipidomic analyses of USS^Hi^ and USS^Lo^ MuSCs. (A)** PCA of the secretomes of USS^Lo^ and USS^Hi^ MuSCs at 1 and 3 dpi. **(B-C)** GO enrichment analysis of downregulated DSPs in USS^Hi^ vs USS^Lo^ at 1 dpi (B) and 3 dpi (C). **(D)** PCA of the lipidomes at 3 dpi. **(E)** Circular bar graph showing the overall Log2(fold change) of total lipid class abundances in USS^Hi^ vs. USS^Lo^ MuSCs.

**Supplementary Figure S4. Single-cell validation of the Ccl2-DTA ablation model. (A)** UMAP projection of captured single cells from both Ctrl and KI groups at 3 dpi. **(B)** Stacked bar chart showing the relative proportions of the identified cell clusters in the Ctrl and KI groups at 3 dpi. **(C)** Violin plots showing canonical marker expression across subclusters. **(D)** UMAP projection colored by USS.

**Supplementary Figure S5. Extended spatial and single-cell transcriptomics analyses of the regenerating niche. (A)** GO enrichment analysis of 8 µm bin data showing up- and downregulated DEGs in KI vs. Ctrl MuSCs **(B)** Spatial expression maps of myogenic markers (*Myod1*, *Myog* and *Myh3*) and immune cells marker (*Ptprc*) in Ctrl and KI muscle sections. **(C)** Dot plot from 8 µm bin data, showing the above marker expression (*Myh3*, *Myog*, *Myod1* and *Ptprc*) in the Injury, Peri-injury and Non-injury zones of Ctrl and KI muscles. **(D)** Annotated cell-type composition across the Injury, Peri-injury and Non-injury zones in Ctrl and KI muscles from nuclei segmentation data. **(E)** High-resolution spatial mapping validating the cell type annotations. The top row displays the specific spatial localization of individual cell populations on representative tissue sections. The bottom row presents spatial feature plots showing the expression levels of corresponding representative marker genes for each cell lineage. **(F)** Dot plot of marker gene expression across spatially annotated cell types. **(G)** Stacked bar chart from analyzing the independent scRNA-seq dataset showing the relative percentage of the annotated cell populations in Ctrl and KI muscles from 1 and 3 dpi. **(H)** Dot plot of marker gene expression across indicated niche cell types in the scRNA-seq dataset. **(I)** CellChat differential interaction strength analysis between Ctrl and KI single-cell transcriptomics data at 1 dpi. MuSC emanated signalings to niche cell populations are highlighted.

**Supplementary Figure S6. FACS gating strategy for EC isolation from wild type mice skeletal muscle at 3 dpi. (A)** FACS gating strategy for CD45- CD31+ Ecs from wild type mice muscle at 3 dpi: FSC-A/SSC-A for bulk cells, FSC-H/FSC-A and SSC-H/SSC-A for doublet exclusion, followed by CD45- (DAPI) and CD31+ (APC) gating.

**Supplementary Figure S7. Baseline MuSC abundance in Ctrl and KI mice. (A)** Flow cytometric quantification of the MuSC fraction (YFP+) in Ctrl and KI muscles at 0 and 1 dpi. About 100,000 cells were sorted by FACS from Ctrl and KI mice muscle at 0 and 1 dpi.
