## Supplementary Figures for "Multiomics Characterization Identifies S1P-secreting USS^High^ Skeletal Muscle Stem Cells as Essential Drivers of Niche Remodeling and Muscle Regeneration"

**Liu X et. al. Suppl. Fig. 1**

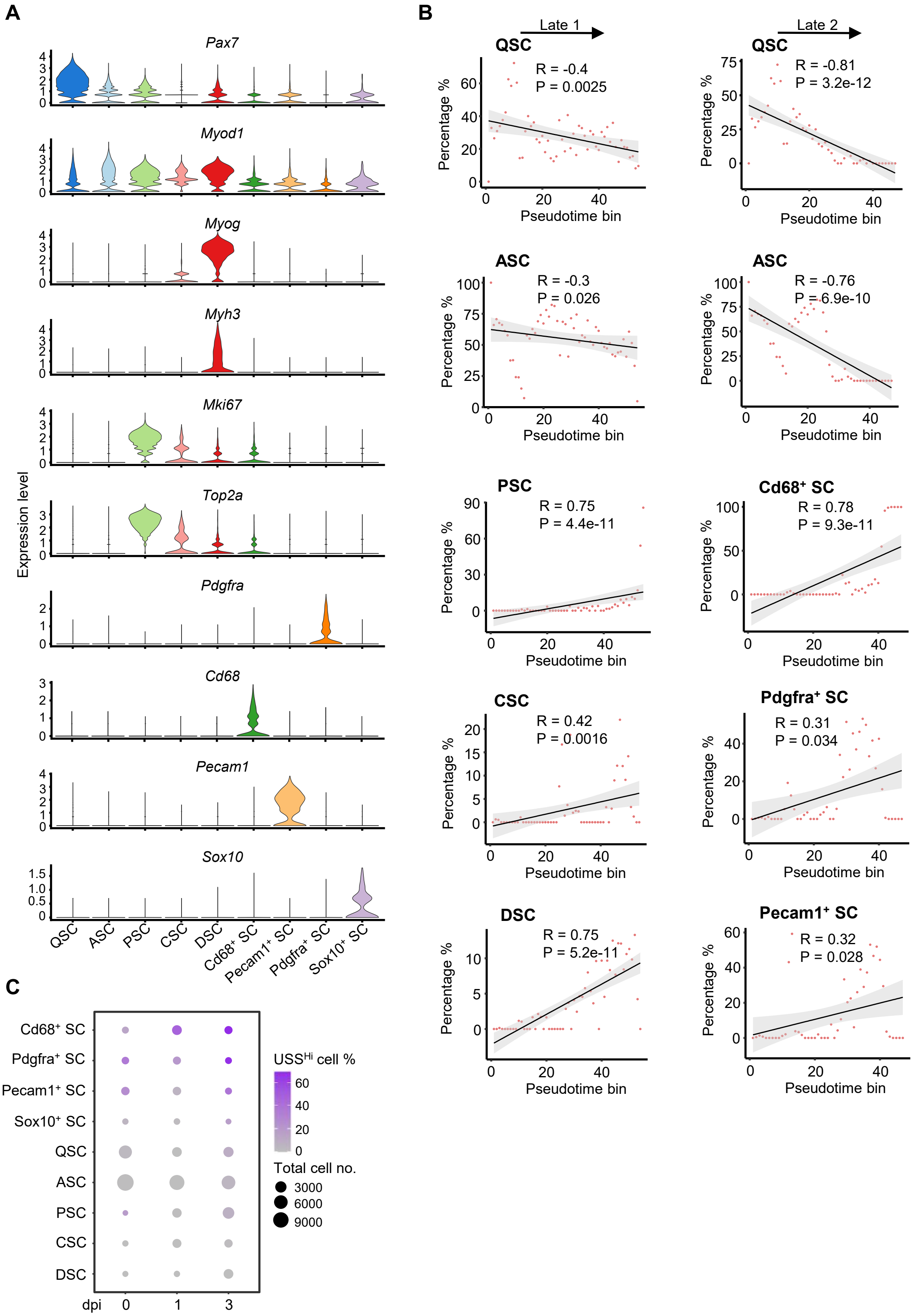

**Liu X et. al. Suppl. Fig. 2**

**A**

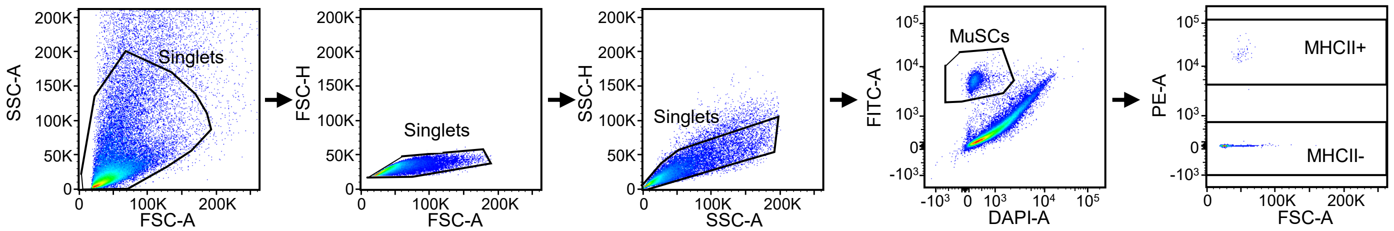

Liu X et. al. Suppl. Fig. 3

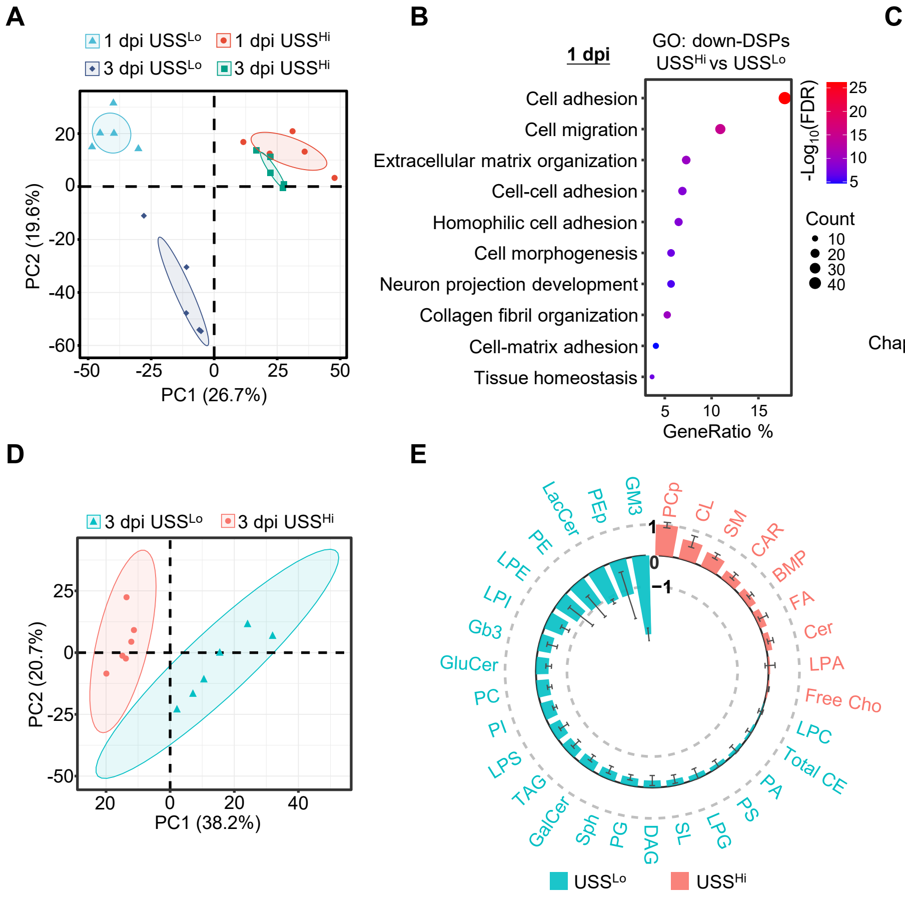

**Liu X et. al. Suppl. Fig. 4**

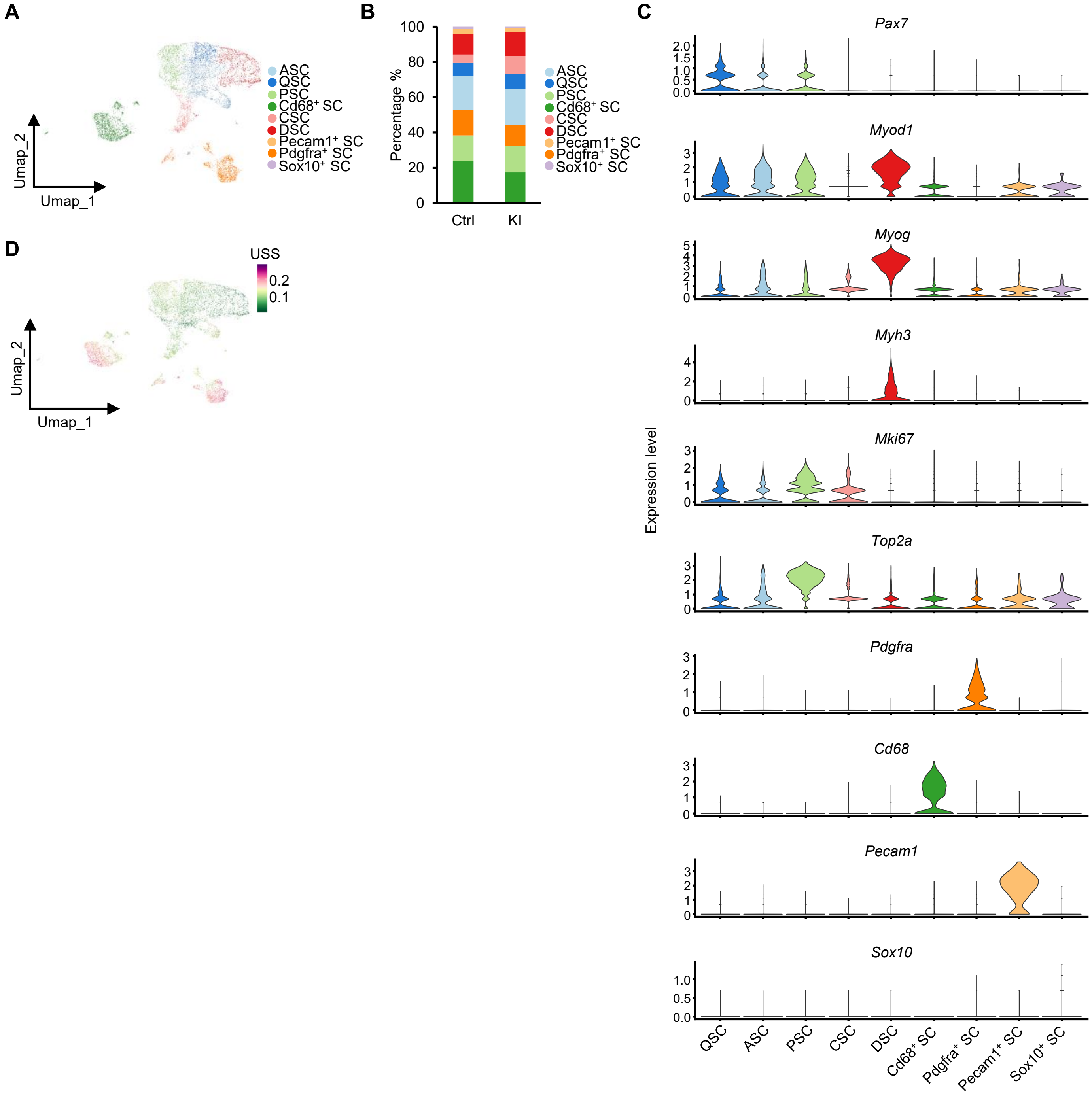

**Liu X et. al. Suppl. Fig. 5**

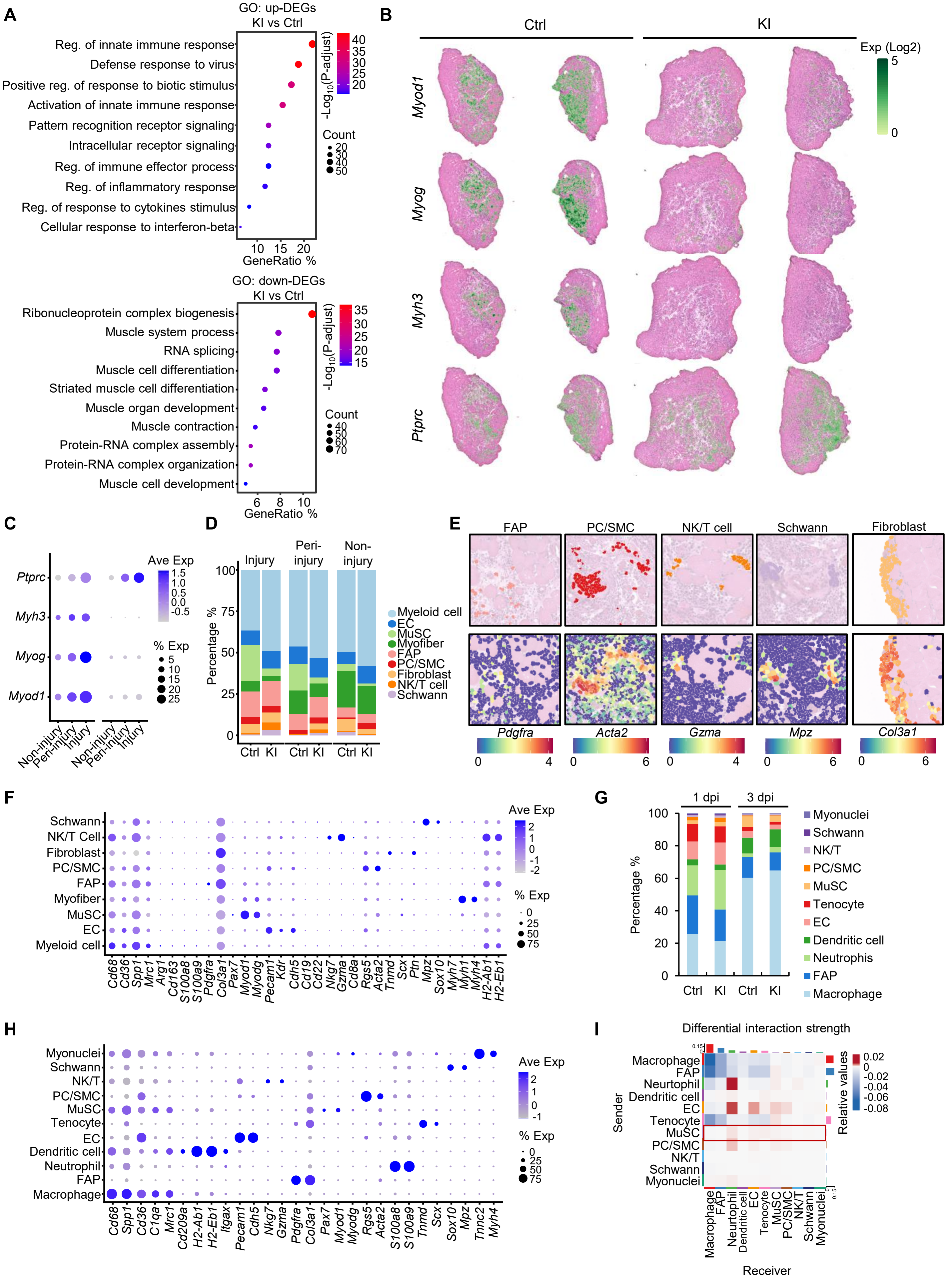

Liu X et. al. Suppl. Fig. 6

**A**

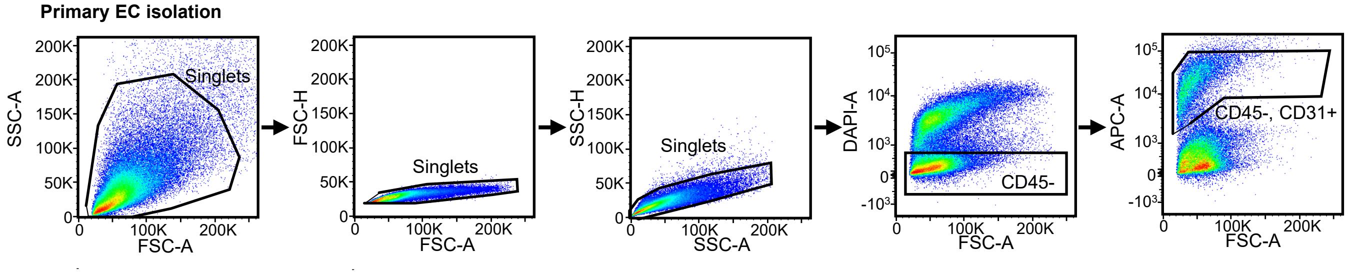

Liu X et. al. Suppl. Fig. 7

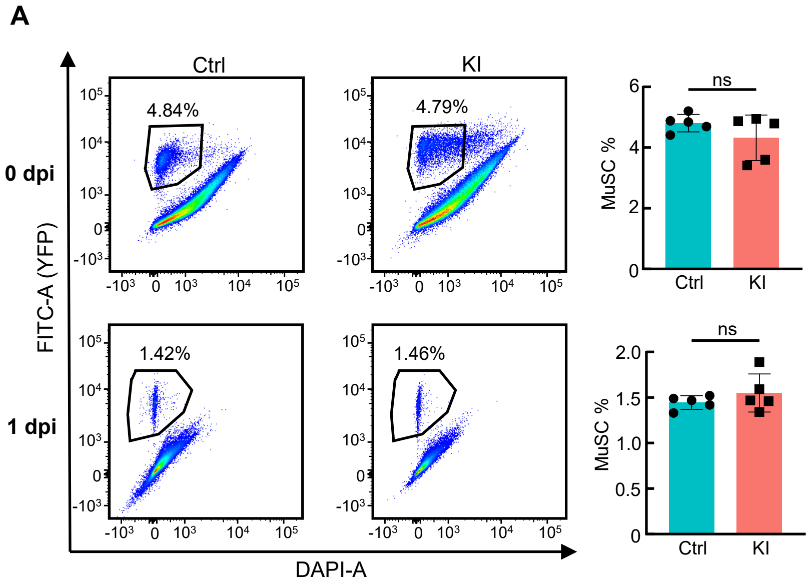
